# Sex-biased genetic programs in liver metabolism and liver fibrosis are controlled by EZH1 and EZH2

**DOI:** 10.1101/577056

**Authors:** Dana Lau-Corona, Woo Kyun Bae, Lothar Hennighausen, David J Waxman

## Abstract

**Background:** Sex differences in the incidence and progression of many liver diseases, including liver fibrosis and hepatocellular carcinoma, are associated with sex-biased expression of hundreds of genes in the liver. This sexual dimorphism is largely determined by the sex-specific pattern of pituitary growth hormone secretion, which controls a transcriptional regulatory network operative in the context of sex-biased chromatin states. Histone H3K27-trimethylation yields a major sex-biased repressive chromatin mark that is specifically deposited by polycomb repressive complex-2, via its homologous catalytic subunits Ezh1 and Ezh2, at many strongly female-biased genes in male mouse liver, but not at male-biased genes in female liver.

**Results:** We used *Ezh1*-knockout mice with a hepatocyte-specific knockout of *Ezh2* to elucidate the sex bias of liver H3K27-trimethylation and its functional role in regulating sex-differences in the liver. Combined hepatic Ezh1/Ezh2 deficiency led to a significant loss of sex-biased gene expression, particularly in male liver, where many female-biased genes increased in expression while male-biased genes showed decreased expression. The associated loss of H3K27me3 marks, and increases in the active enhancer marks H3K27ac and H3K4me1, were also more pronounced in male liver. Many genes linked to liver fibrosis and hepatocellular carcinoma were induced in Ezh1/Ezh2-deficient livers, which may contribute to the increased sensitivity of these mice to hepatotoxin-induced liver pathology.

**Conclusions:** Ezh1/Ezh2-catalyzed H3K27-trimethyation is thus essential for the sex-dependent epigenetic regulation of liver chromatin states controlling phenotypic sex differences in liver metabolism and liver fibrosis, and may be a critical determinant of the sex-bias in liver disease susceptibility.

## Background

Liver disease shows marked sex differences. Hepatocellular carcinoma incidence and mortality is three times higher in men than in women [1, 2], and male mice are more susceptible to chemical-induced hepatic carcinogenesis [3]. Males are also more susceptible to non-alcoholic fatty liver disease, non-alcoholic steatohepatitis and liver fibrosis than females [4–8]. Underlying these sex-biased phenotypical differences are hundreds of genes expressed in liver in a sexually dimorphic manner. This dimorphism is largely regulated by the sex-specific patterns of pituitary secretion of growth hormone (GH), which is intermittent (pulsatile) in males and persistent (near continuous) in females [9]. The sex-specific effects of GH require the GH-activated transcription factor STAT5b [10, 11]. STAT5b cooperates with several sex-biased, GH-responsive transcription factors [12, 13] to regulate sex-specific liver gene expression, working in the context of sex-biased long non-coding RNAs [14, 15], microRNAs [16], and sex-biased chromatin states [17–19]. Trimethylation of histone H3 at lysine 27 (H3K27me3) has been identified as a sex-biased chromatin mark in mouse liver, where a striking male bias in the density of H3K27me3 marks is seen across many of the most highly female-biased genes in male liver, but not at male-biased genes in female liver [17]. Continuous infusion of male mice with GH, which overrides endogenous male plasma GH pulses and imposes a female-like hormonal environment, depletes hepatic H3K27me3 marks at highly female-biased genes in association with induction of female-biased gene expression [19]. These findings suggest that GH regulation of H3K27me3 levels at sex-specific genes contributes functionally to sex differences in liver gene expression and function.

H3K27me3, a hallmark of transcriptional silencing, is deposited by polycomb repressive complex-2 (PRC2), a protein complex involved in cell differentiation, cell-specific identity and cell proliferation [20, 21]. PRC2 has three core components, Suz12, Eed and the homologous catalytic subunits Ezh1 and Ezh2. Ezh1 and Ezh2 both contain a SET domain, which is required for methylation of histone H3 lysine-27. PRC2 can also facilitate transcriptional repression by recruiting protein complexes that recognize H3K27me3 and induce chromatin compaction [22]. Ezh1 and Ezh2 have complementary and compensatory functions and share an overlapping set of target genes [23, 24]. Ezh2 is more active as a methyltransferase than Ezh1 and is preferentially expressed in embryonic and highly proliferative tissues, unlike Ezh1, whose expression persists in adult tissues [23]. Liver size and hepatic progenitor cell expansion are significantly reduced upon deletion of the SET domain of Ezh2 in embryonic mouse liver, which impairs liver differentiation and maturation [25].

PRC2 represses the expression of tumor suppressor genes through both H3K27me3-dependent and H3K27me3-independent mechanisms, promoting tumor formation [26, 27]. Increased levels of Ezh2 and H3K27me3 are found in hepatocellular carcinoma (HCC) and are associated with metastasis and poor prognosis [28]. Ezh2 silences several tumor suppressor miRNAs that are down-regulated in liver cancer [27] and it interacts with highly expressed oncogenic long non-coding RNAs (lncRNAs) to repress target genes in HCC [29, 30]. However, Ezh1 and Ezh2 can also exert anti-tumor effects [31, 32]. Notably, the beneficial effects of Ezh1 and Ezh2 are apparent in adult mouse liver, where the functional loss of both genes induces gene dysregulation accompanied by a severe decrease in liver function, impaired liver regeneration, and induction of liver fibrosis [33], which often leads to development of HCC [34]. Liver steatosis, fibrosis and HCC development are also induced by disruption of GH-STAT5 signaling in mouse liver [35–38], where the metabolic effects of GH signaling loss linked to fatty liver development are more pronounced in males than females [39]. Based on these findings, the sex-dependent pathologies seen in GH signaling-disrupted liver, could, in part, involve the loss of GH-regulated and Ezh1/Ezh2-dependent deposition of H3K27me3 marks required for physiologically balanced expression of sex-biased genes in the liver.

Here, we use an *Ezh1*-knockout mouse model with a hepatocyte-specific knockout of *Ezh2* to investigate the role of H3K27me3 in regulating sex-biased gene expression in mouse liver and the potential impact of this regulation on sex-biased susceptibility to liver disease. Our findings reveal a significant sex-bias in the impact of Ezh1/Ezh2 loss, with a striking preference for depletion of H3K27me3 marks and increased expression of female-biased genes in Ezh1/Ezh2-deficient male liver. Hepatic Ezh1/Ezh2 deficiency is also shown to down regulate many male-biased genes, presumed as a secondary response to the disruption of female-biased gene expression. Finally, we show that many genes associated with liver fibrosis and liver carcinogenesis are differentially responsive to the loss of Ezh1/Ezh2 in male compared to female liver, which may contribute to the observed sex-differences in the incidence and progression of liver cancer.

## Methods

### Animal tissues

Livers from 7-week-old male and female *Ezh1*-knockout mice with a hepatocyte-specific knockout of *Ezh2* (E1/E2-KO mice, also designated Double-knockout (DKO) in the figures and tables) and their age and sex matched floxed littermate controls were generated as described [33]. Briefly, Ezh2^fl/fl^ mice [40] were bred with Alb-Cre transgenic mice [41]; their offspring were then bred with *Ezh1*-knockout mice (Thomas Jenuwein, Research Institute of Molecular Pathology, Vienna, Austria) [42] to generate E1/E2-KO mice. Livers from 2-8 week old male and female CD1 mice (ICR strain) were those described previously [43]. Hypophysectomy and continuous GH infusion of male mice for 14 d using an Alzet osmotic minipump were performed as described [19, 44]. Livers used in this study were obtained from mice housed and handled according to NIH guidelines, and all animal experiments were approved by the Animal Care and Use Committee of National Institute of Diabetes and Digestive and Kidney Diseases.

### qPCR analysis

Liver total RNA (1 μg) was reverse transcribed using the Applied Biosystems High-Capacity cDNA Reverse Transcription Kit (Fisher, Cat#43-688-14). qPCR was performed using Power SYBR green PCR master mix and processed on an ABS 7900HT sequence detection system (Applied Biosystems) or the CFX384 Touch Real-Time PCR detection system (Bio-Rad). For RT-qPCR, raw Ct values were analyzed using the comparative Ct method with normalization to the 18S RNA content of each sample. Primers used for qPCR are shown in Table S1A.

### RNA-seq analysis

Approximately 10% of each liver was snap frozen in liquid nitrogen and used to extract RNA with TRIzol reagent (Invitrogen Life Technologies Inc., Carlsbad, CA). Total liver RNA was isolated from each of 9 individual mouse livers per treatment group (Male (floxed) controls, Female (floxed) controls, Male E1/E2-KO and Female E1/E2-KO). Three RNA-seq libraries (biological replicates) were prepared for each treatment group; each sequencing library was comprised of a pool of RNAs obtained from n=3 individual livers. Sequencing libraries were prepared using the Illumina TruSeq RNA library preparation kit (Illumina, cat# RS-122-2001) and 68 nt single-end sequence reads were obtained on an Illumina HiSeq instrument. RNA-seq data was analyzed using a custom pipeline [44]. Briefly, sequence reads were aligned to mouse genome build mm9 (NCBI 37) using Tophat (version 2.0.13) [45]. FeatureCounts [46] was used to count sequence reads mapping to the union of the exonic regions in all isoforms of a given gene (collapsed exon counting), and differential expression analysis was conducted using the Bioconductor package EdgeR [47]. We identified 11,491 liver-expressed genes, defined as genes expressed at >1 FPKM (Fragments Per Kilobase length of transcript per Million mapped reads) in at least one of the four sex-genotypes analyzed. 1,356 liver-expressed genes were significantly dysregulated in either male or female E1/E2-KO liver [i.e., EdgeR |fold-change| > 1.5 and adjusted p-value (i.e., FDR, false discovery rate) < 0.05 for a comparison of E1/E2-KO male vs control male liver, or for a comparison of E1/E2-KO female vs control female liver]. A set of 1,131 genes showing significant male-female differences in expression in livers of floxed control mice in the E1/E2-KO background strain was identified using cutoff values for sex-differential expression of FDR < 0.01 and FPKM >1; these thresholds empirically corresponded to a >1.2-fold sex-difference in expression (Table S2). A set of 8,021 liver-expressed genes whose expression is stringently sex-independent was defined based on |fold-change| for sex-difference < 1.2 and FDR > 0.1 (Table S3). The responses of these genes to the loss of Ezh1/Ezh2 are shown in Table S2 and Table S3, and are summarized in Table S4.

Differential expression data from livers of male mice treated with GH given as a continuous infusion for 14 d, and for livers of hypophysectomized male and female mice, and their strain and age matched pituitary-intact control livers were obtained from Table S2 and Table S3 of [19] and from Table S3 of [44]. A set of 113 robust female-biased liver-expressed genes was defined as genes with a female/male expression ratio > 2-fold in control mice from each of three different mouse models (Table S5). Raw RNA-seq data from GSE53627 [33] was obtained and re-analyzed using the custom pipeline cited above. Differential expression analysis was performed for the following comparisons: E1/E2-KO males vs. control males (8 months); males treated with CCl_4_ vs. control males; and E1/E2-KO-males treated with CCl_4_ vs. control males (Table S7, Table S8).

For female-biased genes, a percent feminization value was calculated based on each gene’s response to each of the following treatments: Ezh1/Ezh2 loss in male liver; pituitary hormone ablation, as determined by hypophysectomy of male mice; and continuous infusion of male mice with GH for 14 days, as follows:

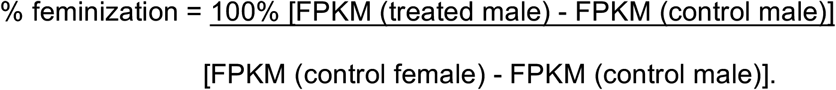

Heat map generation and clustering were carried out using Morpheus (https://software.broadinstitute.org/morpheus/), with average linkage hierarchical clustering implemented on the rows.

### Chromatin preparation and chromatin immunoprecipitation (ChIP)

Chromatin was extracted from frozen liver tissue from each of 6 individual mice per group. Approximately 1 g of frozen liver was submerged in 4 ml of cross-linking buffer [10 mM HEPES (pH 7.6), 25 mM KCl, 0.34 M sucrose, 0.15 mM 2-mercaptoethanol, 2 mM MgCl_2_, and Pierce protease inhibitor (1 tablet per 50 mL of buffer; ThermoFisher Scientific, cat. #A32965)] and homogenized using a glass dounce homogenizer. The homogenate was pushed thorough to a 70-micron cell strainer (Fisher Scientific. #22-363-548) using a 3 ml syringe plunger. The full volume (~ 5 ml) was transferred to a 15 ml conical tube containing 313 μl from a 16% formaldehyde ampule (ThermoFisher Scientific # 28906) mixed with 687 μl of crosslinking buffer, to give a final concentration of 0.83%. The samples were then incubated on a rocker for 5 min at room temperature. Cross-linking was halted by addition of 250 μl of a 2.5 M glycine (pH 8.0) solution (final concentration, 0.1 M), followed by incubation for 5 min at room temperature.

Samples were pelleted (2,500 g for 5 min at 4°C) and washed twice with 10 ml of PBS. Pellets were resuspended in 10 ml of Lysis Buffer 1 [50 mM HEPES (pH 7.5), 140 mM NaCl, 1 mM EDTA, 10% glycerol, 0.5% IGEPAL CA-630 (Sigma-Aldrich, cat. #I8896), 0.25% Triton X-100 (Sigma cat. #T8787) and Pierce protease inhibitor, as above], and incubated on a rocker for 10 min at 4°C. Samples were centrifuged at 2,000 g at 4°C for 5 min, the supernatant was removed, and the pellet was resuspended in 10 ml of Lysis buffer 2 [200 mM NaCl, 1 mM EDTA, 0.5 mM EGTA, 10 mM Tris-HCl (pH 8.0) and Pierce protease inhibitor] and rocked for 5 min at 4°C. Samples were centrifuged at 2,000 g at 4°C for 5 min, the supernatant was removed, and the pellet was resuspended in 2 ml of 1X radioimmunoprecipitation assay (RIPA) buffer [50 mM Tris-HCl (pH 8.1), 150 mM NaCl, 1% IGEPAL CA-630, 0.5% sodium deoxycholate] containing 0.5% SDS and Pierce protease inhibitor. Samples were sonicated for 20 cycles (30 s ON, 30 s OFF) using a Bioruptor Pico sonicator (Diagenode) in 15 ml Diagenode TPX tubes containing 0.3 ml polypropylene beads. A 15 μl aliquot of the sonicated chromatin was incubated at 65°C for 6 hr to reverse cross-links. RNase A (ThermoFisher E053, 10 mg/mL) was added to a final concentration of 0.12 mg/mL and the samples then incubated at 37°C for 30 min. Proteinase K (Bioline BIO-37084, 20 mg/mL) was added to a final concentration of 0.39 mg/mL and samples were digested at 37°C for 2 h. A portion (8 μl) of each sample was analyzed by electrophoresis on a 1% agarose gel to size the DNA fragments, which mostly ranged from 100 to 300 bp. Reversed cross-linked DNA was quantified using a Quant-iT PicoGreen assay kit (Invitrogen). The remaining sonicated chromatin was snap-frozen in liquid nitrogen and stored at −80°C until further use for ChIP. ChIP was performed as reported previously [17, 18] using the following ChIP-validated antibodies: H3K27ac (Abcam cat. # ab4729, 3 μg antibody per 15 μg of sonicated chromatin), H3K4me1 (Abcam cat. # ab8895, 1.2 μg antibody per 15 μg of sonicated chromatin), H3K27me3 (Abcam cat. # ab6002, 2 μg antibody per 10 μg of sonicated chromatin), and normal rabbit IgG (Santa Cruz, cat. # sc-2027, 3 μg antibody per 15 μg of sonicated chromatin). ChIP DNA was quantified using a Quant-iT PicoGreen assay kit (Invitrogen) and analyzed by quantitative PCR (qPCR) using primers that interrogate genomic regions selected as positive controls or as negative controls for each of the histone marks based on our published ChIP-seq data [17].

### ChIP sequencing

Sequencing libraries were prepared for each of the above three histone marks using 20-50 ng of ChIP’d DNA. Libraries were prepared for each of 4 individual livers (biological replicates) for each sex and each genotype (male and female floxed control mice, and male and female E1/E2-KO mice; 16 libraries for each histone mark) using NEBNext Ultra II DNA Library Prep kit for Illumina (NEB, cat. # E7645). NEBNext Multiplex Oligos for Illumina (NEB, Set 1; cat. # E7335, NEB, Set 2; cat. # E7500) were used for multiplexing. The Agencourt AMPure XP system (Beckman Coulter, cat. # A63880) was used for sample and library purification. 50 nt paired-end sequence reads were obtained on an Illumina HiSeq instrument. ChIP-seq analysis was performed using a custom analysis pipeline initially developed for DNase-seq analysis and described elsewhere [48]. Individual biological replicates were validated using standard quality control metrics (FASTQC reports, confirmation of read length, and absence of read strand bias). FASTQ files for validated biological replicates were then concatenated to obtain a single set of combined reads for each condition.

Sequence reads were mapped to the genome using Bowtie 2 (version 2.3.2) [49]. Genomic regions containing a significant number of H3K27me3 reads were identified using SICER (version 1.1, window size 400 bp and gap size 2400 bp) [50] and used for Reads in Peaks Per Million mapped sequence reads (RiPPM) normalization for UCSC browser visualization, as described below. Genomic regions (peaks) enriched for H3K27ac and H3K4me1 sequence reads were discovered using MACS2 (version 2.1.1) [51] with default parameters. ChIP-seq peaks were visualized in the UCSC genome browser (https://genome.ucsc.edu/) after normalization of the genomic regions (i.e., ChIP-seq peak regions) discovered by SICER or MACS2 using RiPPM as a scaling factor, as follows. The peak lists identified for each sample (described above) were merged using mergeBed (BEDtools) to generate a single peak list (peak union). The fraction of reads in the peak union list for each sample was then calculated to obtain a scaling factor. Raw read counts were divided by the per-million scaling factor to obtain RiPPM normalized read counts.

### H3K27me3 differential peak discovery

DiffReps (version 1.55.4) [52] was used to identify genomic regions where H3K27me3 reads showed a significant difference in intensity between conditions being compared (‘differential sites’). These analyses were based on pairwise comparisons of the 4 biological replicates per experimental group for each of the following comparisons: control males vs. control females; E1/E2-KO males vs. control males; E1/E2-KO females vs. control females; and E1/E2-KO males vs. E1/E2-KO females. diffReps differential windows were discovered using the nsd broad option of diffReps using each of four window sizes: 1 kb, 2 kb, 5 kb and 10 kb. For each analysis, the step size was set to 1/10 of the window size. Default statistical testing parameters of diffReps were used: negative binomial test with a p-value cutoff of < 0.0001 for significant windows. Windows with significant differential H3K27me3 marks that were discovered with two or more of the window size settings were consolidated to eliminate redundancy by retaining the diffReps ID number and statistical information for the largest window size setting. Differential windows that were uniquely discovered by any of the four window size settings were also retained. The combined set of retained H3K27me3 differential windows were then filtered by diffReps-determined by FDR <0.05 and |fold-change| > 2 for the experiments groups being compared. The final lists of H3K27me3 differential sites are shown in Tables S6A-S6D.

### H3K27ac and H3K4me1 differential peak discovery

DiffReps (see above) was then applied using the n=4 biological replicate ChIP-seq samples for each chromatin mark, using the 1 kb window setting to identify diffReps differential sites. The differential sites identified were then filtered to retain those sites that overlap a MACS2-identified ChIP-seq peak. The resulting list of retained differential sites was further filtered for downstream analyses by excluding those sites that did not meet the threshold values of diffReps-determined FDR <0.05 and |fold-change| > 2 for the experimental groups being compared. The final lists of H3K27ac differential sites are shown in Tables S6E-S6H, and the final lists of H3K4me1 differential sites are shown in Tables S6I-S6L.

### H3K27me3 peak normalization

Due to the semi-quantitative nature of ChIP-seq methodologies [53], we first identified H3K27me3 regions that are largely unchanged across individuals and genotypes (static H3K27me3 sites). We used diffReps to identify stringent non-differential genomic windows (p-value > 0.1 and |fold-change| < 1.2) in the comparison of control male and E1/E2-KO male livers, and in the comparison of control female and E1/E2-KO female livers. The non-differential H3K27me3 sites identified in male liver were intersected with those identified in female liver, and 1,433 sites with 80% or greater reciprocal overlap across their lengths were retained. Those sites were then filtered by their average raw sequence read counts, and the top 35% of sites (502 sites, >400 average raw sequence reads per site) were retained. Those 502 sites were further filtered to retain the top 75% sites whose H3K27me3 marks were least variant across samples after RPM normalization (376 sites). ChIP-qPCR analysis of a subset of these stringent non-differential H3K27me3 sites (see Fig. 4A, Fig. 4B, below; and Fig. S3, below; primers used for qPCR are shown in Table S1B) confirmed that this approach does indeed identify invariant sites. ChIP-qPCR also revealed moderate compression of the strong differential sites in the ChIP-seq data, as expected. Next, we calculated the fraction of total sequence reads found in these 376 sites for each sample to obtain a scaling factor. Raw sequence read counts were divided by the per-million scaling factor to obtain normalized read counts for each sample. The normalization factor was then used to provisionally override the diffReps normalization results, and thereby obtain a new set of differential sites for each comparison. We observed high overlap (~93%) between the differential sites identified using the standard diffReps parameters, described above, and those identified when using the stringent non-differential site peak-based normalization factors, described in this paragraph. This high overlap validated our decision to use the standard diffReps normalization method to identify H3K27me3 differential sites for all downstream analyses.

### Mapping chromatin marks to genes

We used the output of diffReps to annotate and assign each H3K27me3 differential site (see above) to one of the following categories and to the genes associated with them: ProximalPromoter (site within 0.25 kb of a transcription start site (TSS)), Promoter1k (site within 1 kb of a TSS), Promoter3k (site within 3 kb of a TSS), Genebody (site overlaps the genomic region extending from a gene's promoter to 1 kb downstream of the gene’s transcript end site (TES)), Genedesert (genomic regions that are depleted of genes and are at least 1 megabase long), Pericentromere (region between the boundary of a centromere and the closest gene, excluding the proximal 10 kb of the gene's regulatory region), Subtelomere (defined in a manner similar to pericentromere), and OtherIntergenic (any region that does not belong to any of the above categories) (Table S6A-S6D). H3K27me3 differential sites annotated as ProximalPromoter, Promoter1k, Promoter3k and Genebody, and the genes associated to them, were used for downstream analyses in Fig. 5, below. H3K27ac and H3K4me1 differential sites were mapped to their putative gene targets by GREAT [54] using the following parameters: each RefSeq gene was assigned a basal regulatory domain extending from 5 kb upstream to 1 kb downstream of the TSS, and the regulatory domain was extended in both directions to the nearest gene’s basal regulatory domain up to a maximum of 1,000 kb in one direction [54]. For the results presented in Fig. 6, below, genes that were up-regulated in male E1/E2-KO livers (846 genes) were classified into 8 groups (Table S6M) based on the GREAT gene-mark associations and the overlap of the full H3K27me3 region with the gene body or with the 3 kb genomic region surrounding the gene’s TSS. Of the 846 genes, 226 are stringent sex-independent and 260 are sex-biased (Table S6M). The group classifications for these subsets of genes are shown in Fig. 6C.

### Functional annotation and Pathway Analysis

Differential gene expression data were analyzed using the IPA software suite (https://www.qiagenbioinformatics.com/products/ingenuitypathway-analysis) (QIAGEN Inc). Genes related to liver fibrosis and hepatocellular carcinoma were obtained by searching for the terms “liver fibrosis” and “Hepatocellular carcinoma” in the Diseases and Functions search field. Lists output by IPA were further filtered to exclude chemicals by retaining only terms with an associated Entrez gene ID for mouse. Lists of 217 fibrosis-related genes and 920 hepatocellular carcinoma-related genes used in our analysis are shown in Table S7 and Table S8.

### Statistical analysis

The enrichment for up regulation of sex-biased genes in E1/E2-KO liver as compared to non-sex-biased genes was calculated from the ratio (A/B) divided by (C/D), as shown in this example: A = 240 female-biased genes up-regulated in male E1/E2-KO liver, and B = 842 *minus* 240 = 602 non-female-biased genes up-regulated in male E1/E2-KO liver, where 842 = total number of liver-expressed genes up-regulated in male E1/E2-KO liver; and C = 404 female-biased genes not up-regulated in male E1/E2-KO liver, and D = 10,847 liver-expressed genes not up-regulated in male E1/E2-KO liver (11,491 total liver-expressed genes *minus* 644 total female-biased genes). In this case, (A/B) divided by (C/D) = 10.7-fold enrichment Fisher exact test was used to determine the statistical significance of all the enrichment and depletion calculations (Table S11). Graphical and statistical analyses were performed using GraphPad Prism 7 software. qPCR data are expressed as mean values and either standard errors of the mean or standard deviation for n = 3 to 12 individual mouse livers per group, as specified in each figure legend. Unpaired t-test or one-way analysis of variance (ANOVA) with a Dunnett posttest was used to compare groups to each other, as noted in the figure legends.

### Data availability

All raw and processed RNA-seq and ChIP-seq data for 7-week floxed control and E1/E2-KO mice are available under accession number GSE110934 at Gene Expression Omnibus (https://www.ncbi.nlm.nih.gov/gds/). RNA-seq data for CCl_4_-treated control and E1/E2-KO male mouse livers, and for 8-month control and E1/E2-KO male mouse livers [33] are available under accession number GSE53627.

## Results

### Sex-independent expression of Ezh1 and Ezh2 expression in mouse liver

We examined the expression of *Ezh1* and *Ezh2* in male and female mouse liver from 2 to 8 weeks of age (Fig. 1A). *Ezh1* levels did not change significantly over the postnatal liver developmental period examined. *Ezh2* expression peaked at 2 weeks, when hepatocytes are still proliferating [55], and then progressively declined with age in both sexes. This is consistent with the preferential expression of *Ezh2* in highly proliferative cells and with its decline in postnatal development seen in other mouse tissues [23]. No significant sex-bias in liver expression of either *Ezh1* or *Ezh2* was seen at any of the ages examined. *Ezh1* but not *Ezh2* mRNA levels were greatly reduced in both male and female livers of 7-week-old *Ezh1*-knockout mice with hepatocyte-specific inactivation of *Ezh2* (E1/E2-KO livers) (Fig. 1B, Fig. 1C). However, for both *Ezh* genes, transcription was abolished from the exons encoding the SET domain (Fig. 1B, *red box*), which is essential for histone H3K27 methyltransferase activity [24].

**Fig. 1.**
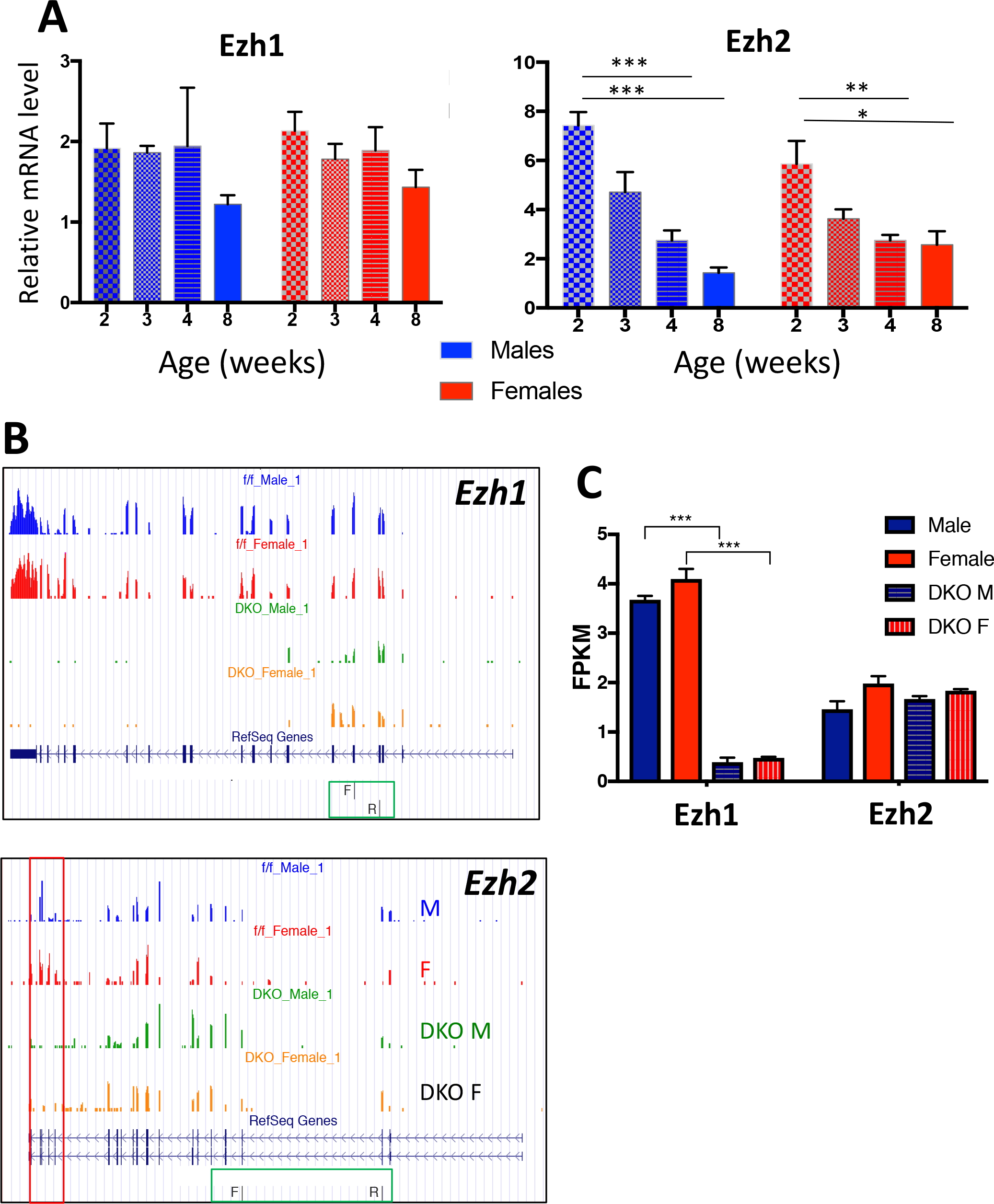
*Ezh1* and *Ezh2* expression in wild-type pre-pubertal and young adult liver and in E1/E2-KO mouse liver. (**A**) Relative expression levels of *Ezh1* and *Ezh2* determined by RT-qPCR in male and female mouse livers at 2, 3, 4 and 8 weeks of age. Data shown are mean ± SEM for n=3 (*Ezh1*) or n=6 (*Ezh2*) individual livers per group. Primers used are shown in Table S1A and their location is marked with a green box in Fig. 1B. Significance values by Student’s t-test are indicated in each figure, as follows: * p < 0.05; ** p < 0.01 and *** p < 0.001. (**B**) UCSC genome browser screenshots of RNA-seq BigWig tracks evidencing the absence in 7-week male and female E1/E2-KO livers of *Ezh1* and *Ezh2* sequence reads from several exons, including those that code for the SET domain (*Ezh1 exons 17-21 and Ezh2* exons 16-19, marked with a *red box*). (**C**) In the case of *Ezh1*, but not *Ezh2*, gene disruption leads to a significant decrease in overall normalized sequence reads, shown in units of FPKM. Data are mean ± SEM values for n=3 individual livers per group. Significance values represent FDR values determined by EdgeR, *** p < 0.001. DKO, Ezh1/Ezh2 double knockout male (M) and female (F) mouse liver.

### Female-biased genes are preferentially de-repressed in Ezh1/Ezh2-deficient male liver

RNA-seq revealed that 1,355 (12%) of 11,491 liver-expressed genes are differentially expressed between E1/E2-KO and control mouse liver. A majority (72%) of the differentially expressed genes were up-regulated in the absence of Ezh1 and Ezh2, as is expected for a genetic deficiency in the capacity for Ezh1/Ezh2-mediated deposition of repressive chromatin marks. More genes showed dysregulated expression in E1/E2-KO males than in E1/E2-KO females (Table S4A). Sex-biased genes comprised 38% of the genes dysregulated in E1/E2-KO liver (435 of 1,131 sex-biased genes; Fig. 2A, Table S4B), and were strongly enriched in both the up-regulated gene set (ES = 9.4, p < 2.2E-16) and the down-regulated gene set (ES = 8.9, p < 2.2E-16) when compared to a set of stringently-sex-independent genes that responded to E1/E2-KO in either sex (Fig. 2A; 413 of 8,021 such genes; Table S4C).

**Fig. 2.**
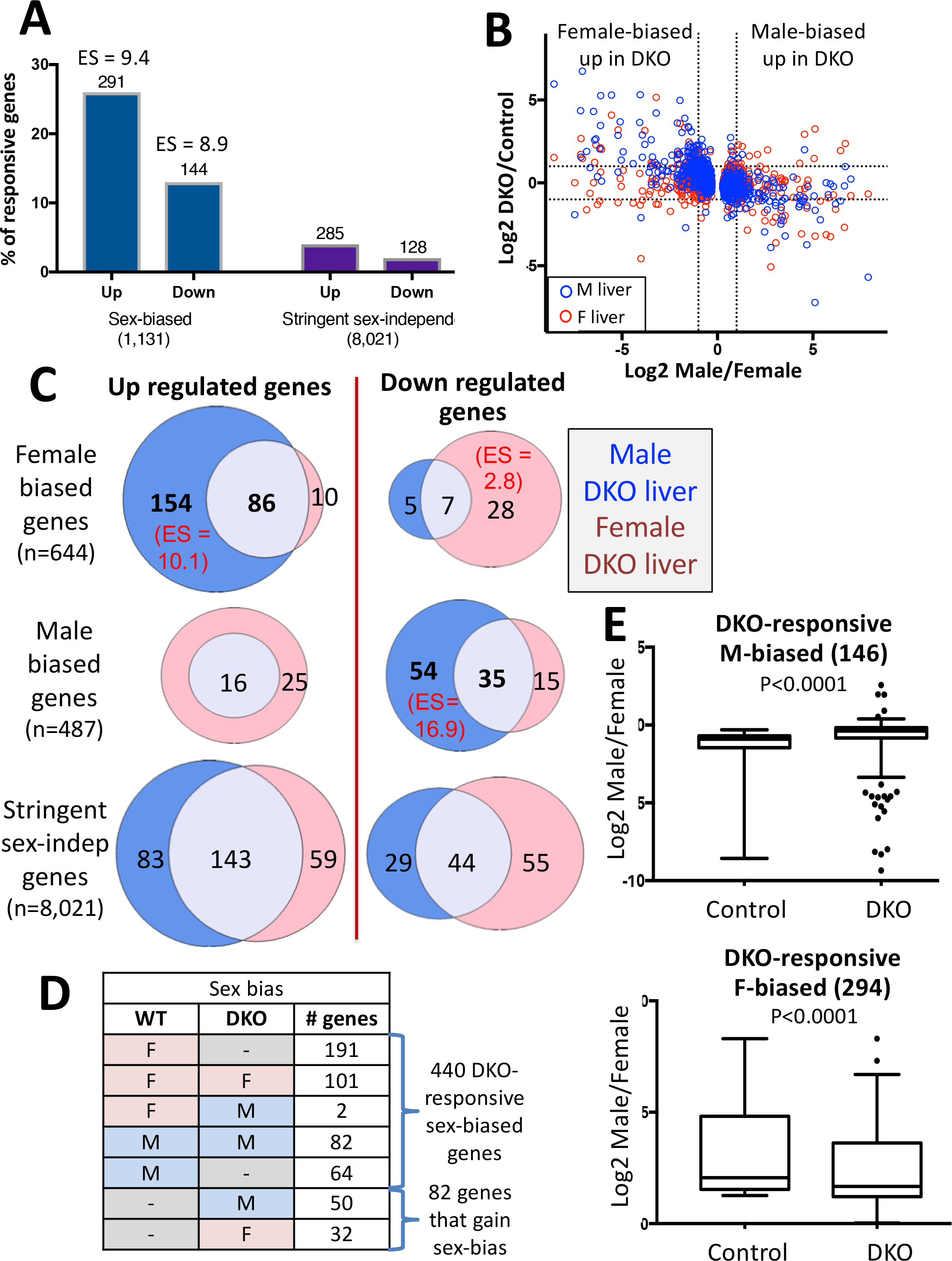
E1/E2-KO de-represses female-biased genes in male liver. (**A**) Percentage of all RefSeq genes that are liver-expressed (FPKM >1) and either sex-biased (male/female FDR < 0.01; n=1,131 genes) (Table S2) or stringently sex-independent (male/female |fold-change| < 1.2 and FDR > 0.1; n=8,021 genes) (Table S3) whose expression in E1/E2-KO liver is significantly changed compared to control liver (|fold-change| >1.5 at FDR <0.05) ('responsive genes'). The number of sex-biased genes that are up or down regulated in E1/E2-KO liver and their enrichment (ES, enrichment score) compared to E1/E2-KO-responsive, stringently-sex independent genes is shown above each bar. (**B**) Log2 expression ratios for E1/E2-KO/control liver vs log2 sex-ratio for 1,131 sex-biased genes, determined in male (blue) and female (red) mouse liver. Dashed lines, significance cutoff values for each comparison. (**C**) Overlap of E1/E2-KO-responsive female-biased genes (and separately, male-biased genes) identified in male liver (blue) vs. female liver (pink). Overlaps are shown separately for genes that are up regulated (*left*) and genes that are down regulated (*right*) in E1/E2-KO liver. Shown at the bottom are the overlaps of stringent sex-independent genes that were E1/E2-KO responsive in male vs. female liver. Significant enrichment scores (ES) compared to a background set of 11,491 liver-expressed genes (FPKM >1) are shown in parenthesis. Five genes responded to the loss of Ezh1/Ezh2 in the opposite direction in male vs. female liver and were excluded from these Venn diagrams: 4 female-biased genes up regulated in male E1/E2-KO and down regulated in female E1/E2-KO, and 1 male-biased gene up regulated in female E1/E2-KO and down regulated in male E1/E2-KO. DKO, Ezh1/Ezh2 double knockout mouse liver. (**D**) Number of E1/E2-KO-responsive sex-biased genes that lose, maintain, or gain sex specificity in either male or female E1/E2-KO mouse liver. Two genes showed a reversal from female to male bias in E1/E2-KO liver and are excluded from this list. Sex bias is either: M, male; F, female; or not sex-biased (dashed line). (**E**) Male/female expression ratios in control liver and in E1/E2-KO liver, for all 146 E1/E2-KO-responsive, male-biased genes (*top*) and for 294 E1/E2-KO-responsive, female-biased genes (*bottom*) (see Table S4). Significance by t-test is as indicated.

H3K27me3 was previously identified as a major sex-biased repressive mark: it is found at many highly female-biased genes in male liver, but not at highly male-biased genes in female liver [17]. Consistent with that finding, many female-biased genes were up-regulated (de-pressed) in E1/E2-KO male liver, while few male-biased genes were up-regulated in E1/E2-KO female liver (Fig. 2B). Further, 154 of 250 up-regulated female-biased genes were exclusively up-regulated in male E1/E2-KO liver, whereas only 10 genes were exclusively up-regulated in female E1/E2-KO liver (Fig. 2C, *top left*). Overall, genes up-regulated in E1/E2-KO male liver were strongly enriched for female-biased genes (ES = 10.1, p < 2.2E-16) compared to all liver-expressed genes. Further, there was a significant enrichment of male-biased genes in the set of genes down-regulated in E1/E2-KO male liver (ES = 16.9, p < 2.2E-16), and a modest enrichment of female-biased genes in the set of down-regulated in E1/E2-KO female liver (ES = 2.8, p = 5.7E-07) (Fig. 2C; also see Table S4). Thus, the loss of Ezh1/Ezh2 preferentially alters sex-biased gene expression in male liver, where many female-biased genes are induced (de-repressed) and male-biased genes are down-regulated. Stringently sex-independent genes did not show a significant sex bias in their response to E1/E2-KO (Fig. 2C, *bottom*).

A majority of all E1/E2-KO-responsive female-biased genes (191 of 294 genes; 65%) lose sex-specificity in the absence of Ezh1/Ezh2 (Fig. 2D), primarily due to their up regulation in male liver. Further, 64 of 146 (44%) E1/E2-KO-responsive male-biased genes lose sex-specificity, primarily due to their down regulation in male E1/E2-KO liver (Fig. 2D; Table S2, column U). The dysregulation of sex-biased genes in E1/E2-KO mouse liver can also be seen by comparing overall gene expression sex ratios in E1/E2-KO liver to control liver (Fig. 2E) and in a heat map (Fig. S1, decrease in color intensity for many genes; column 4 <u>vs.</u> column 1). Whereas the loss of H3K27me3-based repression can directly explain the increased expression of female-biased genes in E1/E2-KO male liver, the decrease in male-biased gene expression is likely a secondary response to E1/E2-KO. This response may involve CUX2, a female-specific repressor of many male-biased genes [12, 56] that is induced 3.7-fold in male E1/E2-KO liver, insofar as 27% of the male-biased genes down-regulated in E1/E2-KO male mouse liver are direct targets of CUX2 (Table S2).

The female-biased genes up-regulated in male E1/E2-KO liver include many cytochromes P450 (*Cyp* genes), sulfotransferases (*Sult* genes) and other drug metabolizing enzymes genes. Interestingly, while female-biased *Cyp2* family members (*Cyp2b9*, *Cyp2c69, Cyp2c40*), were strongly de-repressed in E1/E2-KO male liver, two female-biased *Cyp3* family members were strongly induced in E1/E2-KO female but not E1/E2-KO male liver (*Cyp3a16,* 36-fold increase; *Cyp3a41a*, 4-fold increase), increasing their female-bias in the absence of Ezh1/Ezh2. Several other highly female-biased genes, including *Sult2a1*, *Ntrk2*, *Ptgds*, *A1bg* and *Cyp3a44*, were strongly induced upon loss of Ezh1/Ezh2 in both male and female liver (Table S2, Fig. S2).

### Relationship between GH regulation and Ezh1/Ezh2 repression of female-biased genes in male liver

Hypophysectomy, which ablates circulating pituitary hormones, abolishes ~90% of liver sex differences, and exogenous GH given either in pulses (male plasma GH pattern) or continuously (female-like GH pattern) substantially restores the corresponding sex-specific patterns of liver gene expression [57, 58]. Furthermore, GH given to intact male mice as a continuous infusion feminizes liver gene expression by inducing many female-biased genes and repressing male-biased genes. The induction of female-biased genes by continuous GH is associated with the loss of H3K27me3 marks in male liver, as was shown for four highly female-biased genes [19]. Here, we compared gene responses to Ezh1/Ezh2 loss to gene responses following hypophysectomy [44] or after continuous GH infusion for 14 days in male liver [19] to better understand the relationship between H3K27me3-based repression of female-biased genes and regulation by the sex-specific patterns of GH secretion.

Loss of Ezh1/Ezh2 partially feminized the expression of a subset of strongly female-biased genes in male liver, as exemplified by the strong, albeit incomplete up regulation of *Cyp2b9* – but not *Fmo3* – in male E1/E2-KO liver (Fig. 3A). *Cyp2b9* and *Fmo3* represent two distinct classes of female-biased genes, which were previously defined based on their responses to hypophysectomy. Class I female-biased genes, such as *Fmo3,* require the female, near continuous plasma GH pattern for full expression; consequently, Class I female genes are repressed in female liver by hypophysectomy. In contrast, Class II female-biased genes, such as *Cyp2b9*, are repressed in male liver by the male pituitary hormone profile; consequently, they are de-repressed (i.e., induced) in male liver following hypophysectomy [44, 57]. *Cyp2b9* and *Fmo3* both have strongly male-biased H3K27me3 marks across their gene bodies, which are lost in Ezh1/Ezh2 deficient liver (Fig. 4C). Nevertheless, only *Cyp2b9* is depressed in the absence of Ezh1/Ezh2 (Fig. 3A). The distinct responses of these genes to Ezh1/Ezh2 loss raised the possibility that Ezh1/Ezh2-catalyzed deposition of H3K27me3 marks serves as the underlying mechanism for Class II female-biased gene repression in male liver. However, inconsistent with this proposal, we found that many Class I female-biased genes are also de-repressed in E1/E2-KO male liver, and at a frequency that matches their overall representation in the full set of female-biased genes (Fig. 3B, *top* vs. *bottom*).

**Fig. 3.**
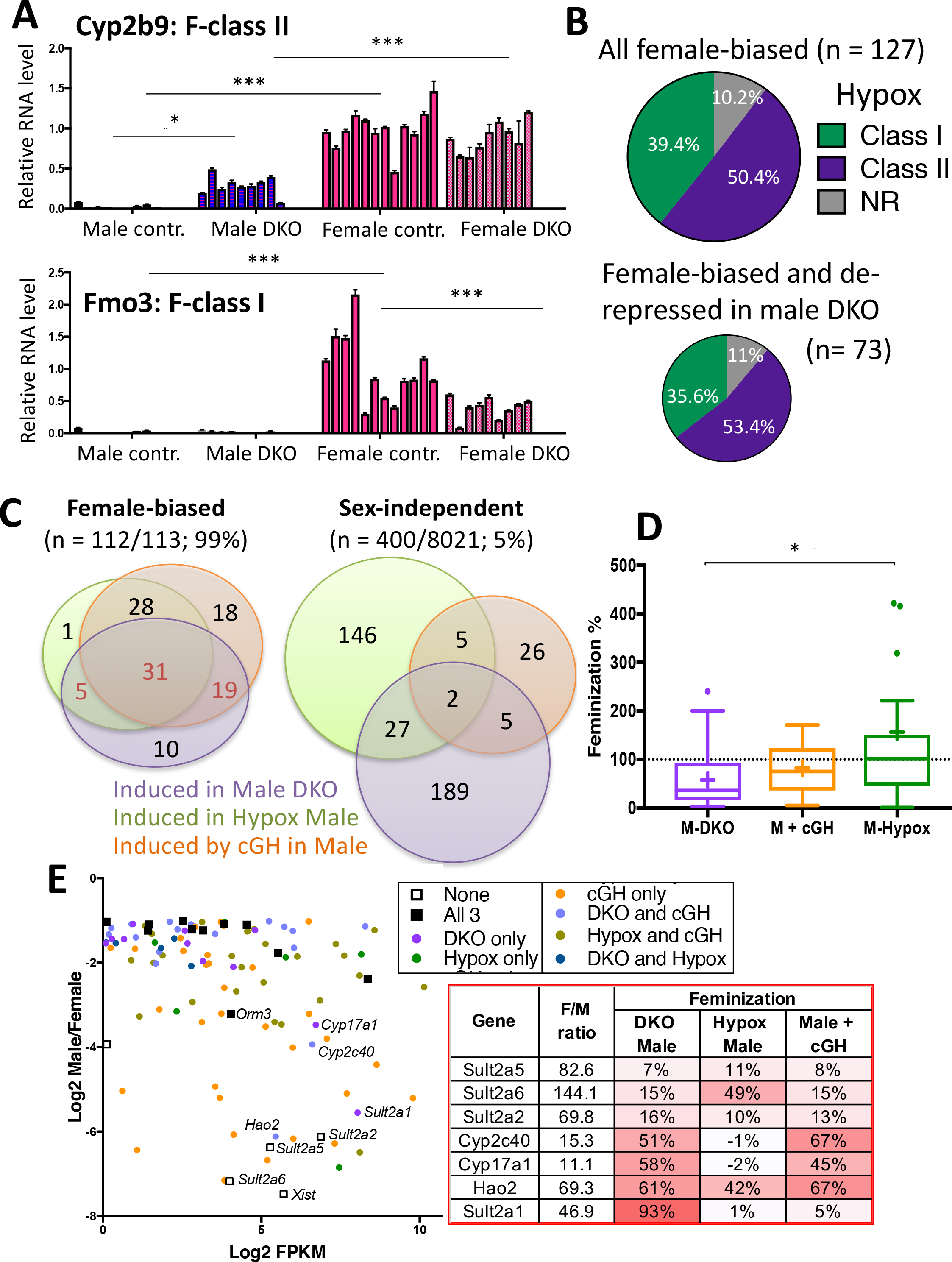
Loss of Ezh1 and Ezh2 partially feminizes the expression of GH-responsive genes. (**A**) Expression of *Fmo3,* a female-biased class-I hypophysectomy (Hypox)-responsive gene (i.e., is down regulated in female liver after Hypox), and *Cyp2b9*, a female-biased class-II Hypox-responsive gene (i.e., is up regulated in male liver after Hypox), in total RNA isolated from floxed control and E1/E2-KO male and female mouse liver. The data shown are mean +/-SD values determined by RT-qPCR for individual livers obtained from 9-12 mice per group. The mean expression value for the control female group was set to 1. Significance values by ANOVA are shown: *, p< 0.05; ***, p< 0.001. Primers used for RT-qPCR analysis are shown in Table S1A. (**B**) Proportion of all female-biased genes (n= 127; female/male expression ratio > 2-fold) and proportion of the subset comprised of 73 female-biased genes that are up regulated in E1/E2-KO male liver and that respond to Hypox and are either class I or class II female-biased genes, or that do not respond to Hypox (NR, not responsive). (**C**) Overlap between female-biased genes (n= 113, male/female expression ratio > 2-fold, EdgeR FDR< 0.01, and FPKM >1 in the sets of control female vs male mouse livers from the three models shown) that are induced: in E1/E2-KO male liver; following Hypox; or after continuous GH infusion for 14 days (cGH). Overlaps are also shown for the set of stringently sex-independent genes, only 5% of genes respond in one of the three mouse models, and where a much greater fraction of the genes that respond to E1/E2-KO do *not* also respond to Hypox or cGH treatment. Expression data for the 113 female-biased genes is presented in Table S5. (**D**) Boxplots showing the distribution of feminization values for 31 female-biased genes induced in male liver in all three mouse models (see panel C, center). Median value, horizontal line in each box; mean value, + sign within or above (green) each box. Statistical significance by ANOVA: *, p < 0.05. (**E**) Graph, in the form of a MA plot, showing log2 male/female ratio vs. gene expression level, in log2 FPKM units, for the above set of 113 female-biased genes. Genes are colored, to indicate which mouse models/mouse treatments result in feminized expression of the gene in male liver by > 50%. Table at the right shows sex differences and percent feminization values for three *Sult2a* family genes largely resistant to feminization and for four other highly female-biased genes that show substantial feminization in E1/E2-KO male liver. The strong feminization of Sult2a6 and Hao2 in hypophysectomized (Hypox) male liver indicates they are class-II female-biased genes. Table S5 shows the percentage feminization values for all 113 genes. DKO, Ezh1/Ezh2 double knockout mouse liver.

To better understand the GH-dependence of E1/E2-KO-responsive female-biased genes, we examined a set 113 robust female-biased genes (Table S5), of which 65 (58%) are up-regulated in E1/E2-KO male liver. We assessed the responses of these 65 genes to two treatments that disrupt normal circulating GH patterns, hypophysectomy and continuous GH infusion. 55 of the 65 genes (85%) were up-regulated in livers of hypophysectomized male mice and/or in livers of male mice infused with GH continuously (Fig. 3C). In contrast, only 15% (34/223) of stringent sex-independent genes induced in E1/E2-KO male liver showed these responses. Further, expression of all but one of the 113 robust female-biased genes (*Xist)* was induced (i.e., feminized) in male liver in one of more of the three models examined (hypophysectomy, continuous GH infusion, and Ezh1/Ezh2-KO) (Fig. 3C, *left*). However, the extent of feminization was substantially lower upon loss of Ezh1/Ezh2 (median feminization, 36%) than following hypophysectomy (median feminization, 102%) (Fig. 3D; Table S5). Moderately female-biased genes were often substantially feminized upon loss of Ezh1/Ezh2 (>50% feminization), whereas the mean feminization was only 29% for highly female-biased genes (F/M > 10). Exceptions include *Hao2, Sult2a1, Cyp2c40 and Cyp17a1* (Fig. 3E). Three highly female-biased sulfotransferases, *Sult2a2, Sult2a5* and *Sult2a6*, showed only partial feminization in all three models (Fig. 3E), despite the loss of Ezh1/Ezh2-dependent H3K27-trimethylation (Fig. 4C, Fig. S2B). Thus, for a subset of female-biased genes, the loss of Ezh1/Ezh2, and thus the capacity to form H3K27me3 repressive marks in male liver, is not sufficient to de-repress gene expression. The repression of these female-biased genes in male liver likely involves mechanisms more complex than simply packaging the gene and its enhancers in H3K27me3 repressive chromatin.

**Fig. 4.**
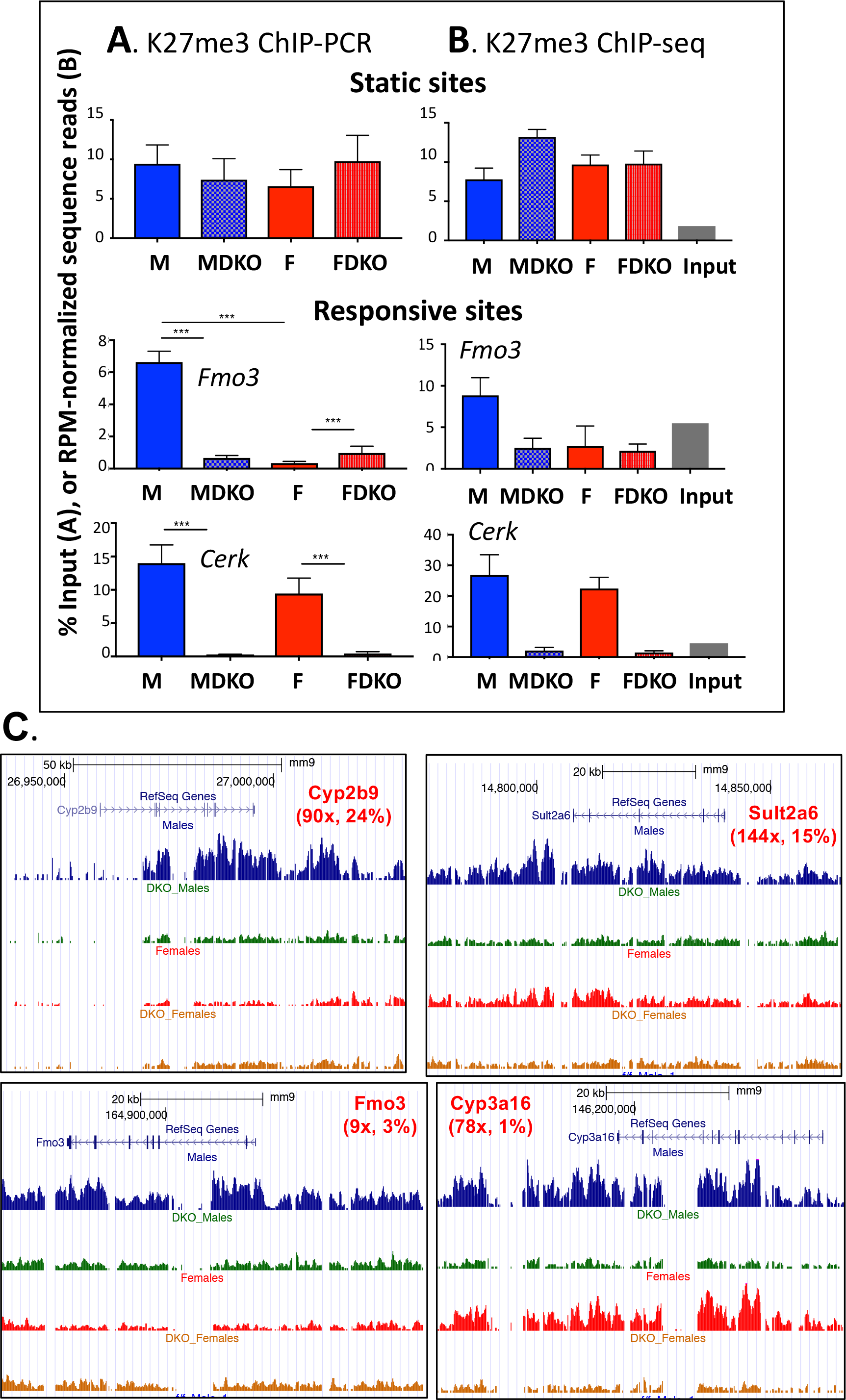
Normalization of H3K27me3 ChIP-seq. ChIP-qPCR validation (**A**) and reads per million (RPM) normalized read counts (**B**) at a K27me3 static site (top row) and at three K27me3 differential sites identified by DiffReps (next three rows). The genomic regions interrogated map to an intergenic static region (qPCR amplicon: Chr 7, 53,631,603-53,631,654), *Hao2* (qPCR amplicon: Chr3, 98,677,644-98,677,901), *Fmo3* (Chr15, 72,993,261-72,993,553) and *Cerk* (Chr1, 164,912,550-164,912,800), as shown. qPCR data shown correspond to % input values for n= 4 individual ChIP DNA samples per group, mean +/-SEM, with significance values determined by ANOVA with Tukey’s multiple comparisons test (***, p< 0.001). B, data are shown for each of the four mouse groups and the input control, as described in Methods. Read counts were obtained for the genomic location corresponding to the qPCR amplicon plus 100 bp, which aims to approximate the 200 bp average sequence library insert size. Primers used for qPCR are shown in Table S1B. DKO, Ezh1/Ezh2 double knockout mouse liver. (**C**) UCSC Browser screenshots showing loss of H3K27me3 sequence reds across the gene bodies of four female-biased genes. The female-bias in H3K27me3 read density in control liver (first and third read tracks) but not in E1/E2-KO liver (DKO; second and fourth read tracks) is also apparent.

### Loss of H3K27me3 marks at E1/E2-KO up-regulated female-biased genes

Global levels of H3K27me3 are reduced in 96% of E1/E2-KO hepatocytes at 3 months of age, without effects on non-parenchymal cells [33]. Here, we investigated the relationship between loss of H3K27me3 marks and the above changes in gene expression. First, to establish the validity of our sequencing results in the absence of a reference epigenome, we identified genomic regions where H3K27me3 marks were significantly lost (differential H3K27me3 sites), as well as regions where the intensity of H3K27me3 marks was unchanged in E1/E2-KO liver compared to control liver (static H3K27me3 sites; see Methods). qPCR analysis of the ChIP’d DNA confirmed that significant changes in H3K27me3 mark intensity occurred at the differential sites but not at the static sites, consistent with the ChIP-seq data for the same sites (Fig. 4A vs. Fig. 4B; Fig. S3). At some differential sites, the loss of H3K27me3 marks in E1/E2-KO liver indicated by sequencing was less complete than indicated by qPCR analysis of the same ChIP’d DNA samples. Thus, the ChIP-seq data underestimates the loss of H3K27me3 at some sites. Nevertheless, we were able to identify several thousand genomic regions with a significant difference in H3K27me3 marks between E1/E2-KO and control liver (Table S6B, Table S6C).

Comparison of control male and female liver identified 538 genomic regions with male-biased H3K27me3 marks vs. only 11 regions with female-biased H3K27me3 marks (Fig. 5A, Table S6A), consistent with the strong male-bias in mouse liver H3K27me3 marks reported previously [17]. Strikingly, this sex bias was abolished in E1/E2-KO liver for all 549 sex-biased H3K27me3 sites; furthermore, 63 other H3K27me3 sites acquired sex-bias in the absence of Ezh1/Ezh2 (Fig. 5A). H3K27me3 marks were decreased at a majority (76%) of the sites dysregulated in E1/E2-KO liver, as expected given the loss of Ezh1/Ezh2. Twice as many H3K27me3 sites were dysregulated in male E1/E2-KO than in female E1/E2-KO livers (2,236 vs 1,096 sites, Fig. 5B), consistent with the greater number of genes dysregulated in male liver (Fig. 2). Sites with H3K27me3 marks down-regulated in both male and female E1/E2-KO livers (male-female common sites, Fig. 5C, Fig. 5D) were enriched for female-biased genes (ES = 2.5, p = 5.59E-04) when compared to a background set of all liver-expressed genes. Female-biased gene enrichment was also seen for sites whose H3K27me3 marks were down-regulated only in male E1/E2-KO liver, and for sites showing down regulation only in female E1/E2-KO liver (Fig. 5E).

**Fig. 5.**
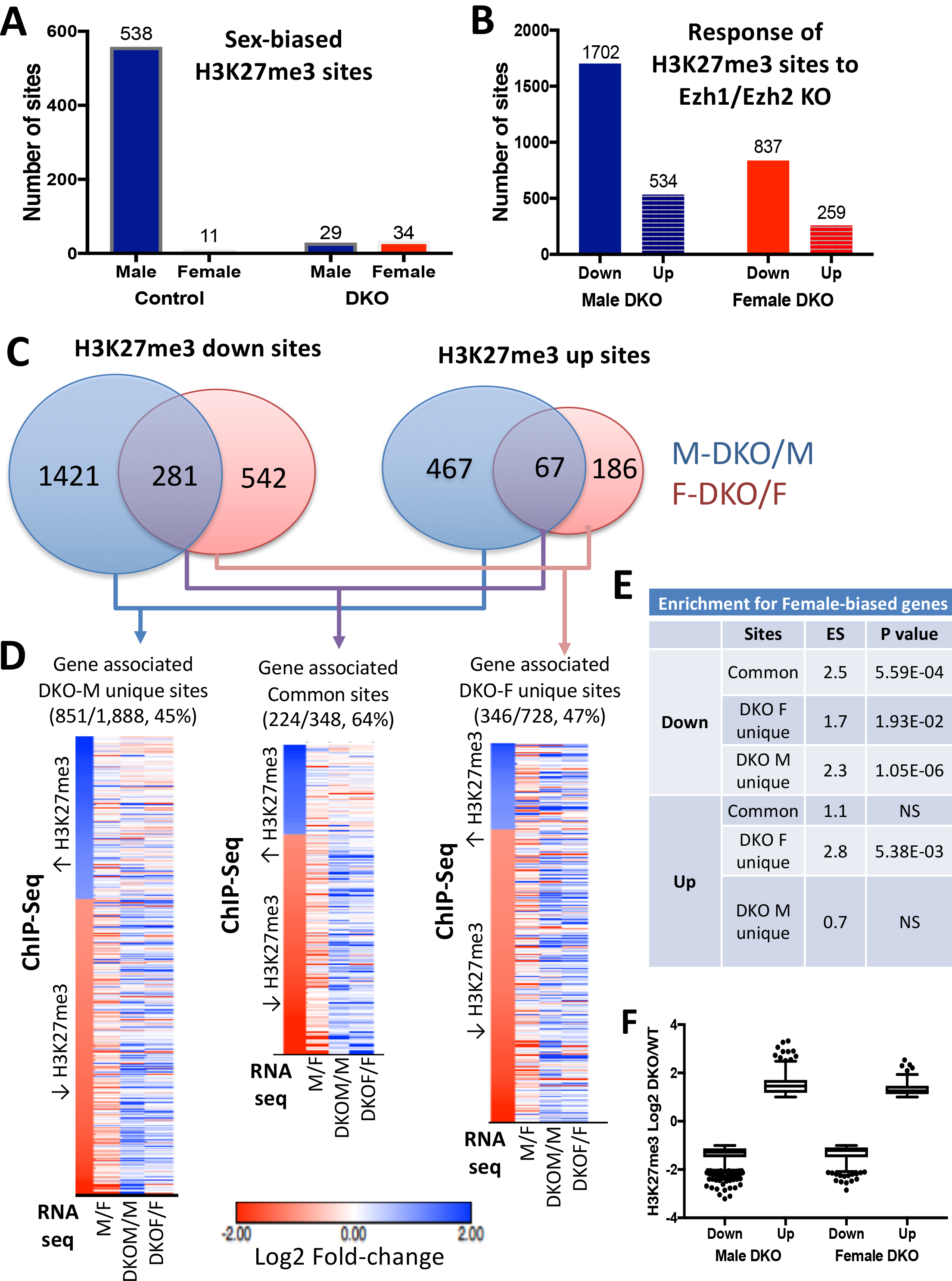
H3K27me3 ChIP-seq. (**A**) Number of sex-biased H3K27me3 sites found in control and in E1/E2-KO male and female mouse liver chromatin, as shown above each column. Sex-bias was lost at all 549 H3K27me3 sites in E1/E2-KO liver. 29 other H3K27me3 sites acquire male bias and 34 sites acquire female bias in the E1/E2-KO livers; the latter sets of sites are associated with 24 genes, of which only 5 are responsive to Ezh1/Ezh2 loss and only one is sex-biased (data not shown). Also see Table S6A. (**B**) Number of H3K27me3 sites that were up regulated or were down regulated in E1/E2-KO male liver, or in E1/E2-KO female liver, when compared to control mice of the same sex, based on Table S6B and Table S6C. (**C**) H3K27me3 sites that are up regulated and down regulated in E1/E2-KO male and E1/E2-KO female liver compared to sex-matched control livers, based on diffReps FDR <0.05, FC >2. Venn diagrams show the overlap between sets of sites for the indicated male-female comparisons. (**D**) Shown on the *left* are heat maps for H3K27me3 sites that are significantly differential only in E1/E2-KO male vs. control male comparison (E1/E2-KO-M unique differential sites) (first column in heat map) and RNA-seq expression ratios for their associated genes (next three columns). Similarly: *Right* heat map, H3K27me3 sites that are differential only in E1/E2-KO female vs. control female comparison (E1/E2-KO-F unique) and their associated genes; and *Middle* heat map, H3K27me3 differential sites common to E1/E2-KO males and E1/E2-KO females and their associated genes. Gene associations for each H3K27me3 site were based on the diffReps tool’s output for those sites located in the gene body or promoter region. Values in the first column of each heat map represent log2 fold-change of the ChIP-seq signal between E1/E2-KO and control liver, and the values for each of the remaining 3 columns represent the log2 fold-change of the gene expression values between the indicated conditions, determined by RNA-seq. Shown above each heat map is the number and percentage of sites that were associated with genes. (**E**) Shown are the enrichment scores and their p-values (see Table S11) for female-biased genes associated with down regulated or up regulated H3K27me3 sites. (**F**) Log2 of E1/E2-KO/control K27me3 signal fold-change at those sites where the density of H3K27me3 marks decreases (Down) or increases (Up) in male and female E1/E2-KO liver. DKO, Ezh1/Ezh2 double knockout mouse liver.

Unexpectedly, H3K27me3 marks were up-regulated in E1/E2-KO liver at ~24% of all H3K27me3 differential sites (Fig. 5B). The extent of up regulation at these sites was similar to the extent of down regulation at other H3K27me3 differential sites (Fig. 5F). H3K27me3 sites up-regulated only in E1/E2-KO-female liver were enriched for female-biased genes (ES = 2.8, p = 5.38E-03; Fig. 5E). The gene targets of the up-regulated H3K27me3 sites (452 genes; gene mapping based on annotations output by diffReps; see Methods) include 64 liver-expressed genes responsive to E1/E2-KO, 25 of which were repressed in either male or female E1/E2-KO liver (FDR < 0.05) (Table S6B, Table S6C). The increase in H3K27me3 marks at these sites could reflect histone mark changes in non-parenchymal cells, where the *Ezh2* gene is intact and presumably still active.

### Ezh1/Ezh2 loss is associated with gain of activating marks

H3K27 can be modified by acetylation to form H3K27ac, an activating mark associated with active enhancers [59]. In the absence of Ezh1/Ezh2, H3K27ac marks can increase and thereby reverse PRC2-mediated gene silencing [24, 60]. ChIP-seq analysis revealed significant increases in H3K27ac and a second activating mark, H3K4me1, at ~900-1,800 sites in male and female E1/E2-KO livers compared to sex-matched control livers; decreases in these activating marks were seen at many fewer (~100-150) sites (Fig. 6A; see Table S6E-S6L for histone mark data). Thus, loss of the capacity to repress chromatin via H3K27-trimethylation is associated with an increase in activating histone marks. Further, whereas male-biased activating chromatin marks (both H3K27ac and H3K4me1) were more than twice as frequent as female-biased sites in control livers, this sex difference was abolished in E1/E2-KO livers (Fig. 6B). The overlap between the sets of sex-biased H3K27ac and H3K4me1 sites in control compared to E1/E2-KO mouse liver was low (Fig. S4), indicating that sex-biased chromatin marks are both gained and lost in Ezh1/Ezh2-deficient liver.

**Fig. 6.**
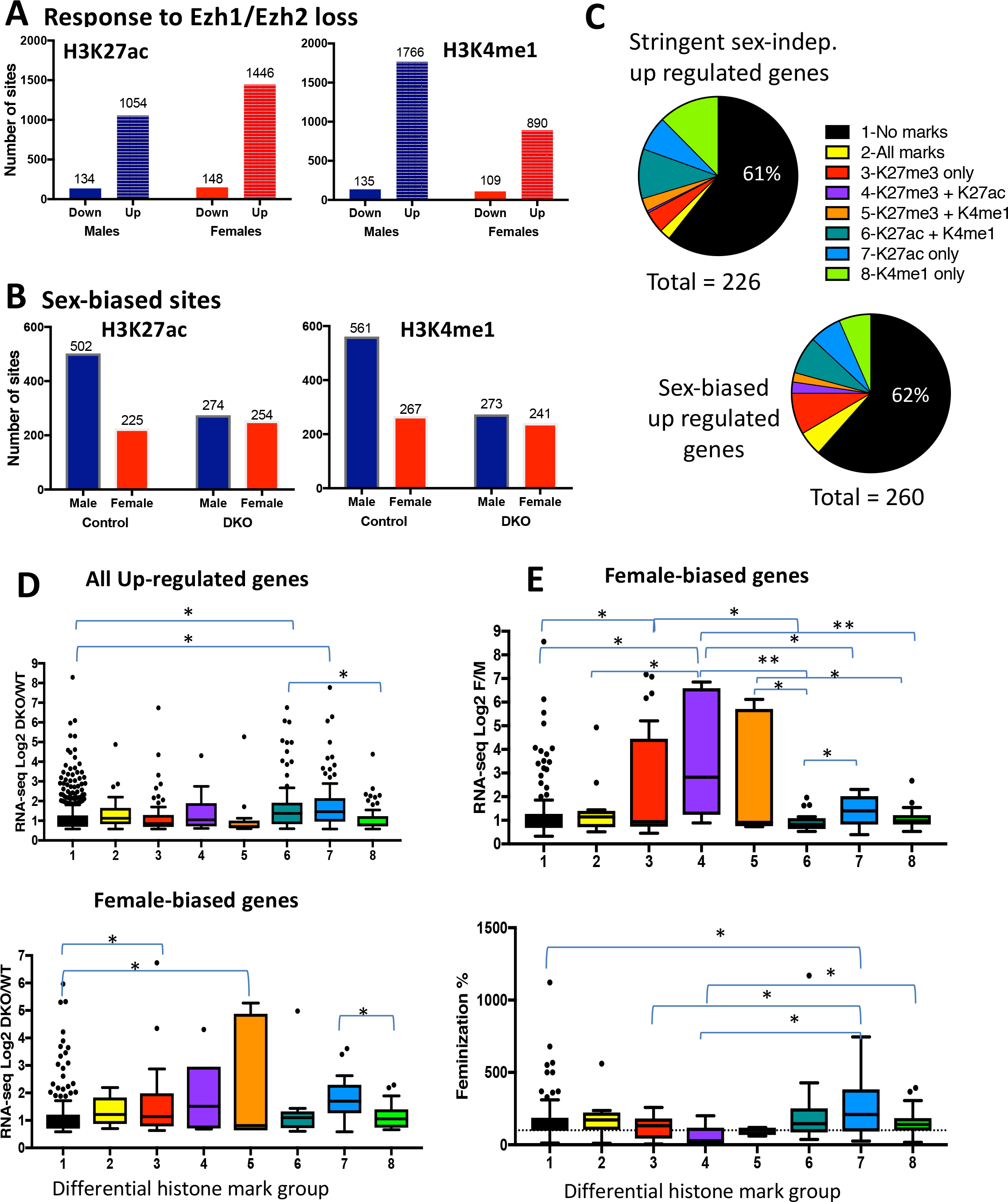
De-repression of gene expression and responsive histone marks. (**A**) Number of sites that show a significant decrease (Down) or a significant increase (Up) in each histone mark in E1/E2-KO male and E1/E2-KO female liver compared to sex-matched control liver. (**B**) Number of male-biased sites (Male) and number of female-biased sites (Female) for H3K27ac and H3K4me1 in control liver, and in E1/E2-KO liver. See listings in Table S6. (**C**) Patterns of histone mark changes associated with E1/E2-KO-responsive genes. *Upper pie chart:* proportion of the 226 stringent-sex independent genes that are up regulated in E1/E2-KO male liver that show a decrease in H3K27me3 marks at their gene body or promoter region, or are associated with an induced H3K27ac or H3K4me1 mark, as determined by GREAT (see Methods). *Lower pie chart:* results shown for the 260 sex-biased genes up regulated in E1/E2-KO male liver. Consistent with data shown in Fig. 2 and Table S4, 846 genes are up-regulated in male liver, of which 260 are sex-biased (154+86+16+4= 260) and 226 are sex independent (143 + 83 =226). (**D**) *Upper* boxplots show the distribution of log2 fold-change values of E1/E2-KO-male/control male for the 846 genes up-regulated in male E1/E2-KO liver in each of the groups, based on their association with responsive histone marks. *Lower* boxplots show the distribution of log2 fold-change values of E1/E2-KO-male/control male only for the 244 female-biased genes up regulated in male E1/E2-KO liver, in each of the groups based on their association with responsive histone marks. **(E)** *Upper* boxplots show the distribution of log2 sex-ratio for the 244 female-biased genes up regulated in male E1/E2-KO liver, in each of the groups based on their association with responsive histone marks. *Lower* boxplots show the distribution of feminization percentages for the 244 female-biased genes up-regulated in the male E1/E2-KO livers, in each of the groups based on their association with responsive histone marks. Significance values by t-test are indicated in each figure as follows: *, p < 0.05; **, p < 0.01; DKO, Ezh1/Ezh2 double knockout mouse liver.

We mapped H3K27ac and H3K4me1 differential sites to their putative target genes using GREAT [54], and H3K27me3 differential sites were assigned to a gene if they overlapped the gene body or 3 kb surrounding its TSS. 846 genes up-regulated in E1/E2-KO males compared to control male liver were classified into 8 groups based on the patterns of differential histone marks associated with each gene (Table S6M). The distribution of histone mark patterns across the 8 groups was generally similar for sex-biased genes as for stringent sex-independent genes up-regulated in E1/E2-KO male livers (Fig. 6C, *top* vs. *bottom*), although groups 2-4, which have differential H3K27me3 marks, were more frequent in the sex-biased gene set, and groups 6 and 8, which have differential H3K4me1 marks but not differential K27me3 marks, were more frequent in the sex-independent gene set. A majority (~60%) of the genes up-regulated in E1/E2-KO male liver were not associated with any differential histone marks (group 1; Fig. 6C). The up regulation of these genes in the absence of changes in H3K27me3, H3K27ac or H3K4me1 marks could be due to de-repression of their transcriptional activators. For example, 13% of the sex-biased genes in group 1 (no differential chromatin marks) are direct targets of the female-specific activator of female-biased genes CUX2, whose expression increases in Ezh1/Ezh2-KO male liver in association with loss of H3K27me3 and increased H3K27ac and H3K4me1 marks (Table S6M). Genes associated with increases in H3K27ac marks, either with or without induction of K4me1 marks (groups 6 and 7), showed significantly greater up regulation than genes without any differential marks (group 1) (Fig. 6D, top). For female-biased genes, loss of K27me3 with or without induction of K4me1 (groups 3 and 5) resulted in greater induction of gene expression than having no associated differential marks (group 1).

Female-biased genes de-repressed in male E1/E2-KO liver that showed both a loss of H3K27me3 and a gain of H3K27ac marks had a significantly higher sex-bias than genes in other differential mark groups (Fig. 6E, top, group 4 vs. groups 1,2,6,7,8). Highly female-biased genes (female/male expression ratio > 10) in group 1 (no differential marks) includes *Sult2a4*, which although it is induced by > 60-fold in E1/E2-KO male liver, only reaches 17% of the expression level of control female liver. However, this group also includes *Cyp2c40* and *Cyp17a1,* whose expression was significantly feminized in E1/E2-KO male liver (51% and 58%, respectively) (Table S6M). Genes showing a gain in H3K27ac marks alone showed greater feminization than genes having no differential marks or loss of K27me3 alone (Fig. 6E, bottom; group 7 vs. groups 1,3,4). Finally, 10 of 12 female-biased genes showing a loss of H3K27me3 marks and an increase in both H3K27ac and H3K4me1 marks (group 2) were fully feminized in male E1/E2-KO liver.

### Ezh1/Ezh2-dependent, sex-differential regulation of liver fibrosis genes and HCC-related genes

Male mice and humans are more susceptible to liver fibrosis than females [4–8] and show a male predominant incidence and progression of HCC [1, 2, 61]. Ezh1 and Ezh2 can contribute to the onset and progression of liver fibrosis, as male E1/E2-KO livers acquire a nodular appearance with portal and periportal inflammation and collagen deposition by 8 months of age, along with substantial impairment of liver function [33]. Further, E1/E2-KO male liver shows increased susceptibility to the fibrogenic and hepatotoxic effects of carbon tetrachloride (CCl_4_) [33]. Strikingly, almost half of all liver fibrosis-associated genes (104 of 217 genes, 48%) and HCC-related genes (425 of 920 genes, 46%) were significantly changed in expression (FDR < 0.05, |fold-change| >2) in male E1/E2-KO compared to control liver at either 7 weeks or 8 months of age, or in livers of E1/E2-KO or control livers of male mice exposed to a regimen of CCl_4_ that induces hepatotoxicity and liver fibrosis (Fig. 7A; Table S7, Table S8).

**Fig. 7.**
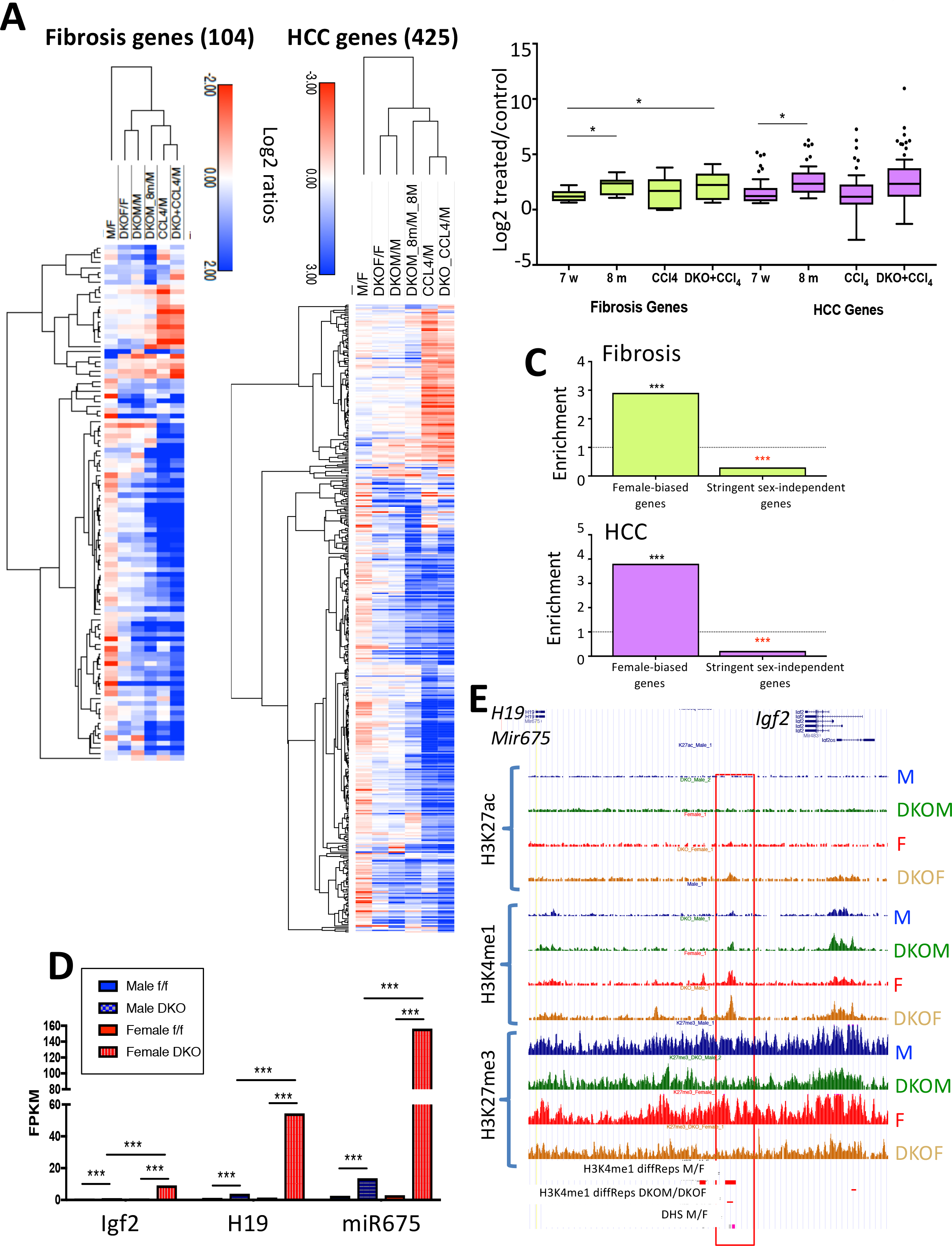
Sex-biased dysregulation of liver fibrosis and HCC related genes. (**A**) Heat maps showing log2 expression ratios for 104 responsive fibrosis-related genes and for 425 responsive HCC-related genes for the 6 indicated comparisons: control male vs. control female, E1/E2-KO female vs. control female, E1/E2-KO male vs. control male, 8-month-old E1/E2-KO male vs. 8-month-old control males, males treated with CCl_4_ and E1/E2-KO males treated with CCl_4_ compared to their age-matched controls. Color bars for ratios ranging ±2 or ±3 log2, as indicated. Average linkage hierarchical clustering implemented on the rows. (**B**) Boxplots showing the distribution of log2 expression changes for 9 fibrosis-related genes (green) and 52 HCC-related genes (purple) that are up regulated in both 7-week and 8-month E1/E2-KO male liver. Values shown for 7-week E1/E2-KO males, 8-month E1/E2-KO males, males treated with CCl_4_ and E1/E2-KO males treated with CCl_4_ compared to their age-matched controls, as described in Methods. Statistical significance obtained by Student’s t-test and indicated as follows: * P < 0.05; ** P < 0.01 and *** P < 0.001. The nine fibrosis-related genes are listed in table S7 and the 52 HCC-related genes are listed in Table S8 (**C**) Enrichment, or depletion, of female-biased genes and stringent sex-independent genes for being in the set of dysregulated fibrosis-related or HCC related genes as compared to all liver expressed genes. (**D**) Expression of *Igf2, H19* and *miR675* determined by RNA-seq (FPKM values) in 7-week male and female control livers and in livers of male and female E1/E2-KO mice. Data shown are mean expression levels based on n=4 individual livers per group. Significance values are based on FDR values determined by EdgeR: ** P<0.01, and *** P < 0.001. (**E**) UCSC genome browser screenshot of the *Igf2*-*H19*-*Mir675* gene locus. Shown are normalized sequence read tracks for H3K27ac, H3K4me1 and H3K27me3 ChIP-seq data. This gene locus has a female-biased DNase hypersensitive site [63] flanked by a female-biased H3K4me1 site that is further induced in E1/E2-KO female liver (horizontal red bars at the bottom), consistent with the greater gene induction seen in female liver. DKO, Ezh1/Ezh2 double knockout mouse liver.

Similar numbers of fibrosis and HCC-related genes were induced in E1/E2-KO male livers at 7 weeks of age as at 8 months (Table 1), even though no overt liver histopathological changes were apparent at 3 months of age [33]. A majority of the dysregulated genes were up-regulated (Table 1) and there was a significant increase in the degree of induction with advancing age (Fig. 7B). Livers of E1/E2-KO male mice exposed to CCl_4_ showed the greatest number of responsive fibrosis and HCC-related genes, and the highest degree of up regulation (Table 1, Fig. 7B), consistent with a previous conclusion based on an analysis of a smaller number of genes [33]. Moreover, the dysregulated sets of 104 liver fibrosis-related genes and 425 HCC-related genes were significantly enriched for female-biased genes and depleted of stringent sex-independent genes when compared to all liver-expressed genes (Fig. 7C). More fibrosis and HCC-related genes were dysregulated in E1/E2-KO female compared to E1/E2-KO male liver (Table 1), consistent with the male bias in liver disease susceptibility.

**Table 1.**
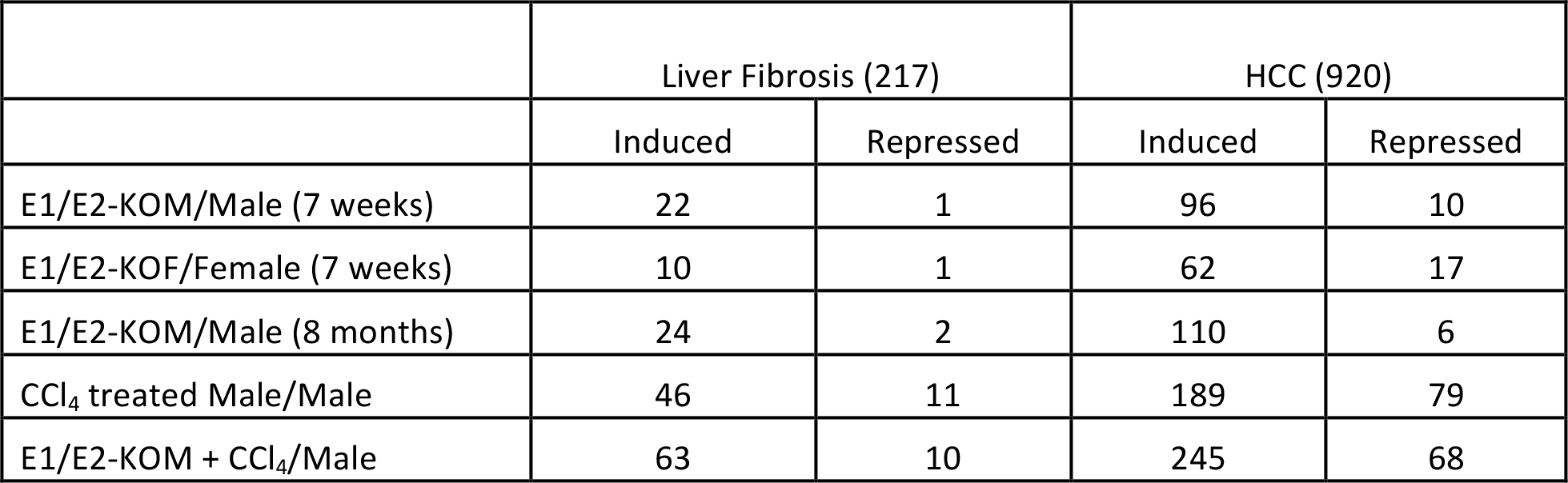
Number of fibrosis-related or HCC-related genes induced or repressed in E1/E2-KO livers or with each of the indicated treatments.

We also identified 32 sex-independent genes that are more highly up-regulated in E1/E2-KO female than E1/E2-KO male liver, and thus acquire female-biased expression in the absence of Ezh1/Ezh2 (Fig. 2D, Table S9). Three genes that show the most significant up regulation are *Igf2*, the lncRNA gene *H19*, and the microRNA *miR675* (Fig. 7D). *Igf2* and *H19* are imprinted genes associated with HCC development [29, 62]. The *Igf2-H19* locus has a female-biased DNase hypersensitive site, identified previously [63], and a female-biased H3K4me1 mark that increases in female E1/E2-KO liver (Fig. 7E). Thus, while there is moderate loss of H3K27me3 marks in both sexes, female-biased increases in activating marks may explain the strong induction of these genes seen in female liver (Fig. 7E).

## Discussion

Ezh1 and Ezh2 are epigenetic modifiers that catalyze H3K27-trimethylation essential for liver homeostasis and regeneration. Loss of Ezh1 and Ezh2 in hepatocytes leads to liver fibrosis, impaired liver function and increased susceptibility to the hepatotoxic effects of CCl_4_ [33]. Marked sex differences characterize the incidence, progression and severity of these liver pathologies, however, the underlying molecular basis for these sex differences in liver disease is only partially understood [64]. Our previous work identified H3K27me3-based gene repression as a sex-biased epigenetic regulatory mechanism in mouse liver [17], suggesting that sex differences in Ezh1/Ezh2-catalyzed deposition of H3K27me3 marks may contribute to the striking differences in liver pathophysiology between the sexes. Here, we used male and female Ezh1/Ezh2 double knockout (E1/E2-KO) mice to investigate the role of Ezh1 and Ezh2 in the regulation of sex-biased genes in mouse liver, and to discover any sex-dependent effects of Ezh1/Ezh2 loss on genes associated with liver disease. We found that hepatic Ezh1/Ezh2 deficiency induces a strong, preferential dysregulation of sex-biased, as compared to sex-independent, genes. Notably, many female-biased genes were significantly de-repressed in E1/E2-KO male liver, in association with the loss of H3K27me3 marks across the gene body, while comparatively few male-biased genes were correspondingly de-repressed in E1/E2-KO female liver. Rather, many male-biased genes were down-regulated in E1/E2-KO male liver, which likely is a secondary response to the up regulation of female-biased genes expression. Thus, Ezh1/Ezh2-based repression of female-biased genes is a major epigenetic regulatory mechanism controlling sex-biased gene expression in male mouse liver. We also found that Ezh1/Ezh2 deficiency up regulates many genes associated with liver fibrosis and HCC in male liver, and that these changes are seen by 7 weeks of age, which precedes the histopathological changes seen in 8-month-old mice [33]. Finally, we found that liver fibrosis- and HCC-associated genes are differentially responsive to the loss of Ezh1/Ezh2 in male compared to female liver, which may contribute to the sex differences in disease incidence and progression.

Sex differences in liver gene expression are primarily regulated by sex-specific patterns of pituitary GH secretion. GH secretion is intermittent in males, whereas in females, pituitary GH release is more frequent, resulting in persistent stimulation of GH signaling in hepatocytes. GH-responsive liver transcription factors, including STAT5b and the STAT5-dependent repressors BCL6 [65] and CUX2 [12], are key mediators of the sex-dependent transcriptional actions of GH and operate in the context of GH-regulated sex differences in chromatin accessibility [63] and sex-biased chromatin states [17]. H3K27me3 is a strikingly sex-biased epigenetic regulatory factor that is specifically associated with strong repression of highly female-biased genes in male liver [17]. Consistently, female-biased genes are significantly enriched in the gene set up-regulated in E1/E2-KO male liver (Fig. 2). However, the feminization of gene expression upon loss of Ezh1/Ezh2 was, in many cases, only partial. This contrasts with the more complete feminization achieved in two other mouse models that we examined, namely, continuous infusion of GH in male mice, which overrides the male, pulsatile plasma GH pattern and induces a majority of female-biased genes within 7 days [19], and ablation of pituitary hormone secretion by hypophysectomy, which de-represses Class II female-biased genes in male liver [44]. Thus, while Ezh1/Ezh2-catalyzed H3K27-trimethylation may repress female-biased genes in male liver, the loss of H3K27me3 marks alone is generally not sufficient for full gene activation, and in some cases, is largely ineffective. Thus, *Fmo3*, a highly female-specific gene (female/male liver expression ratio = 78) was not induced in E1/E2-deficient male liver, despite the extensive loss of male-biased H3K27me3 marks across the gene body (Fig. S2D). One possibility is that gene de-repression is dependent on distal enhancers, which may be subject to distinct epigenetic regulatory mechanisms. Another female-biased gene, *Cyp17a1* (female/male expression ratio = 11-fold), was substantially de-repressed in male E1/E2-KO liver (58% feminization) but showed an unexpected increase, rather than a decrease, in H3K27me3 marks in both male and female E1/E2-KO liver (Fig. S2C).

The de-repression of female-biased gene expression in E1/E2-KO male liver was in many cases accompanied by increases in the active enhancer marks H3K27ac and H3K4me1. H3K27 acetylation cannot occur on nucleosomes where H3K27 is already trimethylated by Ezh1/Ezh2. H3K27ac marks prevent PRC2 binding, antagonize repression by PRC2 [24], and are often enriched in the absence of PRC2 [66]. These findings indicate that the increased expression of female-biased genes in E1/E2-KO male liver is likely a direct result of de-repression caused by the loss of H3K27me3 marks and the subsequent gain in H3K27ac and other activating chromatin marks. Indeed, an increase in activating marks (H3K4me1 and H3K27ac) was associated with stronger induction of gene expression (Fig. 6D top, group 6 and group 7 vs group 1).

Our findings highlight the role of Ezh1/Ezh2-based repression of female-biased genes in male liver as an important mechanism to enforce liver sex differences. 37% of female-biased genes were significantly de-repressed in E1/E2-KO male liver (Fig. 2C), indicating that Ezh1/Ezh2 is responsible – either directly or indirectly – for a substantial fraction of the epigenetic control of female-biased genes. Further, the actions of Ezh1/Ezh2 are sex-biased, with many more genes dysregulated and more widespread loss of sites of H3K27-trimethylation occurring in male than in female liver. The absence of a sex bias in liver Ezh1/Ezh2 expression (Fig. 1) indicates that other factors control the sex differential activity of Ezh1/Ezh2 in adult mouse liver. Indeed, this repression is controlled by circulating GH patterns, whose continuous infusion in male mice induces loss of H3K27me3 marks at female-biased genes in association with their widespread de-repression in male liver [19]. Little is known about the molecular mechanisms by which PRC2 complex and its Ezh1/Ezh2 catalysts are recruited to their specific chromatin targets, in general, and specifically in this case, how plasma GH patterns regulate the sex-dependent interactions between PRC2 and its female-specific gene targets. PRC2 physically associates with several long non-coding RNAs (lncRNAs) [24], which may contribute to the target gene specificity of PRC2’s actions. Further studies are needed to determine whether any of the ~200 liver-expressed, nuclear lncRNAs that show sex-biased and GH-regulated expression in mouse liver [15] contribute to the recruitment of PRC2 to female-biased genes repressed by Ezh1/Ezh2 in male liver.

Several highly female-biased sulfotransferase genes, including *Sult2a2, Sult2a5* and *Sult2a6*, failed to be substantially feminized in Ezh1/Ezh2-deficient male liver. Hepatic expression of these genes was also only partially feminized in male mice when the male, pulsatile pattern of GH stimulation is abolished by hypophysectomy, or when circulating GH profiles are feminized in continuous GH-infused male mice. (Fig. 3). The partial feminization achieved in the latter two mouse models, where GH signaling is disrupted, could be due to early, irreversible postnatal effects of GH, which may imprint (program) liver gene expression patterns [67]. In the case of *Sult2a5* and *Sult2a6,* the high levels of H3K27me3 across the gene body in wild-type male mouse liver were largely abolished in E1/E2-KO male liver (Fig. S2B). Nevertheless, expression of these *Sult2a* genes only reached 7 to 16% of their normal, wild-type female level of expression. Further study is needed to elucidate the mechanisms that establish early, irreversible epigenetic differences in male liver, which may include GH-regulated DNA methylation of gene regulatory regions [68].

E1/E2-KO male mice develop liver fibrosis at 8 months of age, and at 3 months of age, when histopathological abnormalities are not yet apparent, they show much greater susceptibility to the hepatotoxic effects of CCl_4_ as compared to control mice [33]. Here, we found that genes associated with liver fibrosis and HCC are significantly up-regulated in livers of male E1/E2-KO mice in an age-dependent manner and following exposure to the hepatotoxin CCl_4_ (Fig. 7), which correlates with the severity of the liver phenotype [33]. E1/E2-KO female liver showed up regulation of the fewest number of fibrosis/HCC-related genes, consistent with the slower disease progression seen in female liver [69]. Moreover, H3K27me3 marks decreased and/or activating histone marks increased at 50% of the fibrosis- and HCC-related genes up-regulated in 7-week-old E1/E2-KO male liver, similar to the full set of E1/E2-KO up-regulated genes (Table S6M).

Female-biased genes dominate the set of fibrosis/HCC-related genes that were up-regulated in E1/E2-KO male liver. This raises the question of why increased expression of these female-biased genes leads to an increase in liver fibrosis, in particular in male Ezh1/Ezh2-deficient mice [33]; whereas, in wild-type liver, the higher expression of these genes compared to male liver is associated with decreased susceptibility to liver fibrosis and liver disease. The answer may relate to our observation that the set of female-biased genes up-regulated in Ezh1/Ezh2-deficient male liver includes genes that confer protection from HCC in wild-type female liver. One example is *Hao2,* which is down-regulated in HCC, and its expression inversely correlates with metastasis and survival [70]. *Hao2* is strongly induced in E1/E2-KO male liver, but to only ~60% the level of control (wild-type) female liver, and this increase in expression may not be not sufficient to counteract the severe liver injury generated by loss of Ezh1/Ezh2. Furthermore, other female-biased genes that have been recognized as tumor suppressors in HCC, such as *Trim24* [71, 72], are not up-regulated in male E1/E2-KO liver. Further studies are needed to determine the extent to which the higher expression of such liver disease-protective genes in female liver contributes to sex-differences in liver fibrosis and liver disease. Studies such as these take on added significance, given efforts to use Ezh2 inhibitors for treatment of HCC [73].

We identified sex-independent HCC-related genes that were more strongly up-regulated in female than in male E1/E2-KO liver, and thereby acquire female specificity in E1/E2-KO liver. These genes include *Igf2*, *H19* and *miR675* (Fig. 7E, Table S10). *Igf2* and *H19* are adjacent, imprinted genes [74] that show aberrant imprinting and epigenetic abnormalities in HCC [29]. *H19* is highly expressed in proliferative tissues, including liver regeneration after injury [74], and its first exon encodes miR-675, whose overexpression promotes liver cancer [62]. The strong up regulation of these three genes in female E1/E2-KO livers suggests E1/E2-KO female mice may show increased susceptibility to HCC compared to wild-type female liver in response to CCl_4_ and other hepatotoxins.

In addition to extensive up regulation of gene expression following loss of H3K27me3 repressive marks, we identified a significant number of genes that were down-regulated in the absence of Ezh1 and Ezh2, in both male and female liver. In some cases, down regulation was associated with an unanticipated increase in gene proximal H3K27me3 marks (Fig. 5D). While H3K27me3 is generally regarded as a repressive mark, H3K27me3 is enriched at the TSS of bivalent genes, and its enrichment at promoters is associated with active transcription [75]. The partial loss of H3K27me3 marks at E1/E2-KO-responsive genes seen here could represent signal originating from non-parenchymal cells in the liver, where Ezh1/Ezh2 expression is unchanged [33]. Further, as the loss of Ezh1/Ezh2 increases cell proliferation [76], residual H3K27me3 marks could perhaps be explained by the presence of immature hepatocytes, where the Alb-Cre transgene is not yet active [41].

The greater incidence of HCC in males has been attributed to the antagonistic effects of the androgen receptor (AR) and estrogen receptor-α (ERα) activation on hepatocyte proliferation and liver gene expression. Moreover, estrogen has been shown to have a protective role in HCC by reducing the production of inflammatory mediators and by its effects on the hypothalamic-pituitary-gonadal axis including the modulation of GH secretion [1]. AR and ERα mediated transcriptional regulation is dependent on FOXA1 and FOXA2 pioneer factors [77]. This study provides more evidence on the importance of chromatin dynamics in the sex-biased regulation of liver pathophysiology.

## Conclusions

Ezh1/Ezh2-dependent, H3K27me3-based repression is essential to establish and maintain GH-regulated sex differences in liver gene expression. Loss of Ezh1 and Ezh2 in hepatocytes preferentially de-represses the expression of many female-biased genes in male mouse liver in association with the loss of H3K27me3 marks and the acquisition of activating histone marks. Finally, males and females show significant differences in the regulation of liver fibrosis and HCC-related genes by Ezh1/Ezh2, which may contribute to the sex bias in liver disease progression.

## List of Abbreviations

ChIP: chromatin immunoprecipitation
E1/E2-KO mice, or DKO (double-knockout): *Ezh1*-knockout mice with a hepatocyte-specific knockout of *Ezh2*
ES: enrichment score
FC: fold-change
FDR: false discovery rate
FPKM: fragments per kilobase per million mapped sequence reads
GH: growth hormone
H3K27me3: histone-H3 trimethyl-lysine 27
HCC: hepatocellular carcinoma
PRC2: polycomb repressive complex-2
RiPPM: Reads in Peaks Per Million mapped sequence reads

## Declarations

### Ethics approval and consent to participate

Not applicable

### Consent for publication

Not applicable

### Availability of data and material

The datasets generated and/or analyzed during the current study are included in this published article and its supplementary information files. Raw and processed sequencing files generated in this study are available at GEO under accession number GSE110934.

## Authors’ contributions

DL-C and DJW conceived and designed the study. WKB and LH generated the E1/E2-KO mice and provided liver tissues for analysis. DL-C carried out all of the laboratory experiments and data analyses. DL-C and DJW jointly wrote and edited the manuscript for publication.

## Acknowledgements

We thank Andy Rampersaud of the Waxman Lab for developing the sequencing analysis pipeline used in this study.

## Supplemental Tables listing

**Table S1**. qPCR primers used mRNA analysis and for validation of H3K27me3 static and H3K27me3 differential sites.

**Table S2**. 1,131 liver-expressed genes that showed a significant sex-bias in expression in control (floxed) mouse liver.

**Table S3.** 8,021 liver-expressed genes that show stringently sex-independent expression.

**Table S4A.** Responses of all liver-expressed genes (11,491 genes; FPKM >1) to loss of Ezh1/Ezh2 in male and female E1/E2-KO liver.

**Table S4B.** Responses of sex-biased genes to loss of Ezh1/Ezh2 in male and female E1/E2-KO liver.

**Table S4C.** Responses of stringently sex-independent genes to loss of Ezh1/Ezh2 in male and female E1/E2-KO liver.

**Table S5**. Set of 113 robust female-biased genes and their responses to E1/E2-KO, hypophysectomy, and continuous GH infusion in adult male liver.

**Table S6A-S6D.** Differential H3K27me3 sites in autosomes identified by diffReps.

**Table S6E-S6H.** Differential H3K27ac sites in autosomes identified by diffReps.

**Table S6I-S6L.** Differential H3K4me1 sites in autosomes identified by diffReps.

**Table S6M.** Up-regulated genes associated with differential histone marks.

**Table S7**. Expression data for a set of 217 genes involved in liver fibrosis.

**Table S8**. Expression data for a set of 920 genes associated with hepatocellular carcinoma (HCC).

**Table S9.** Listing of 82 genes that gain sex-bias in E1/E2-KO mouse liver.

**Table S10.** Imprinted genes that gain sex-bias in the E1/E2-KO.

**Table S11.** Contingency tables for the enrichments shown in Fig. 2, Fig. 5, and Fig. 7.

---

**Fig. S1.**
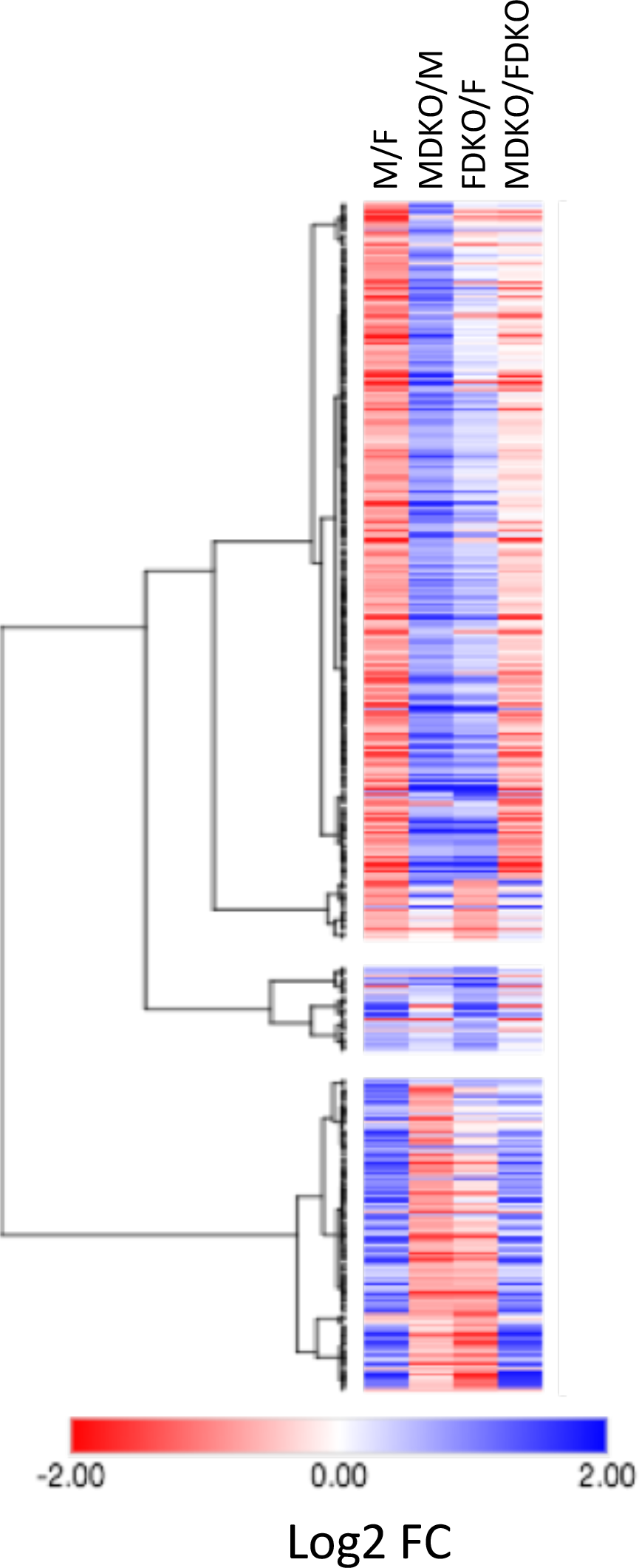
Heat map for the set of 440 E1/E2-KO-responsive sex-biased genes, based on log2 fold-change values for each of the four indicated comparisons. DKO, Ezh1/Ezh2 double knockout mouse liver.

**Fig. S2.**
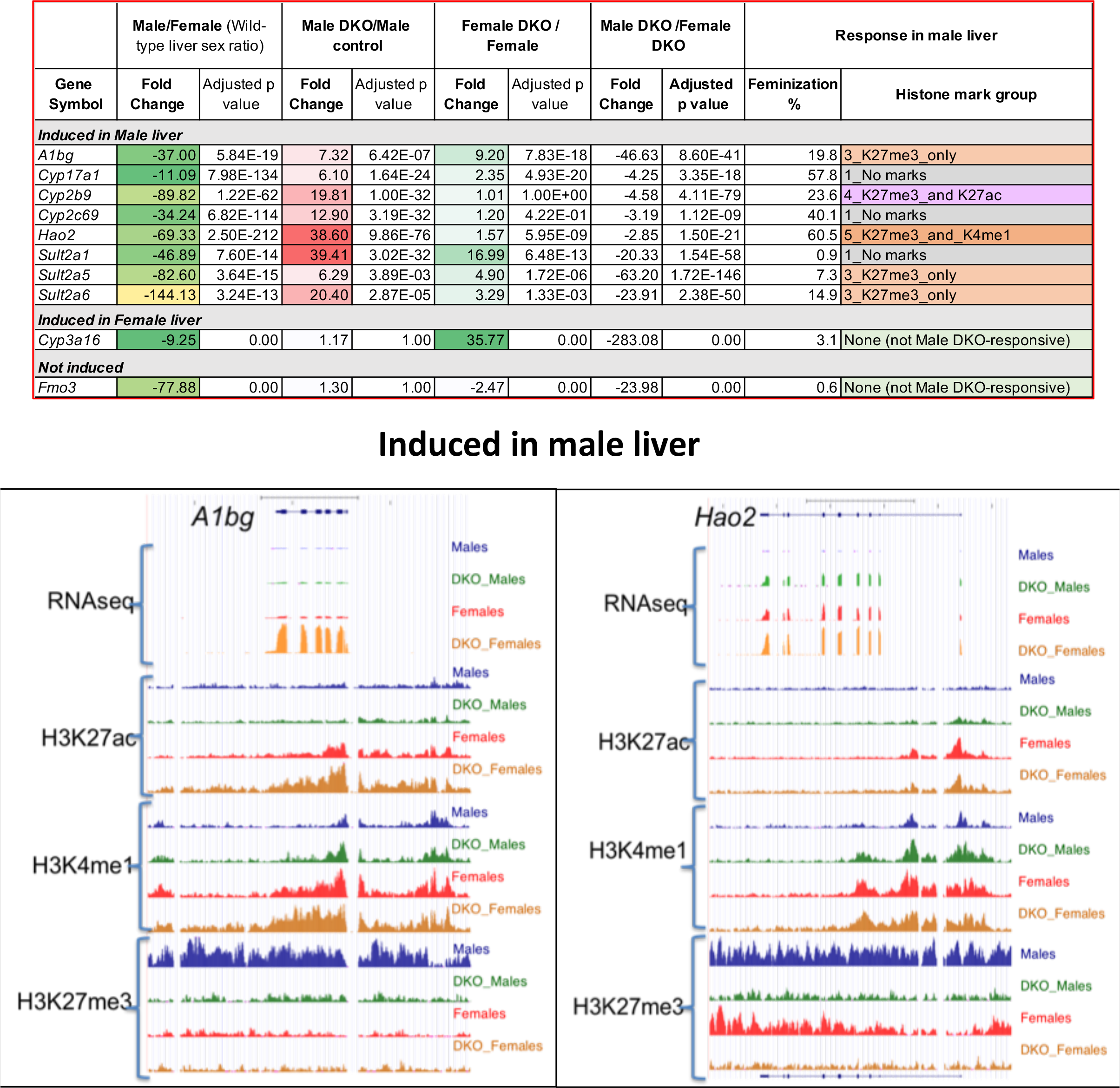
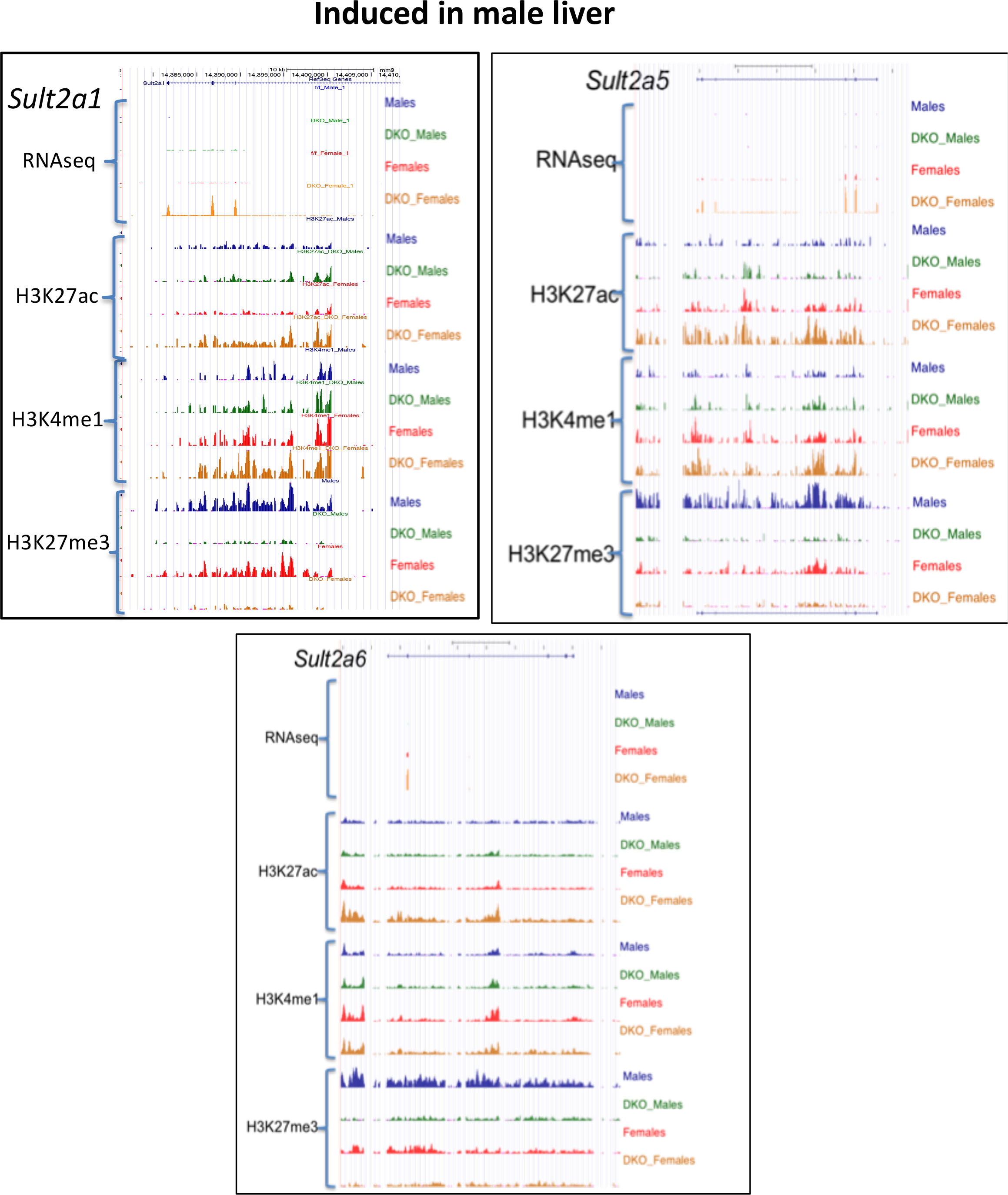
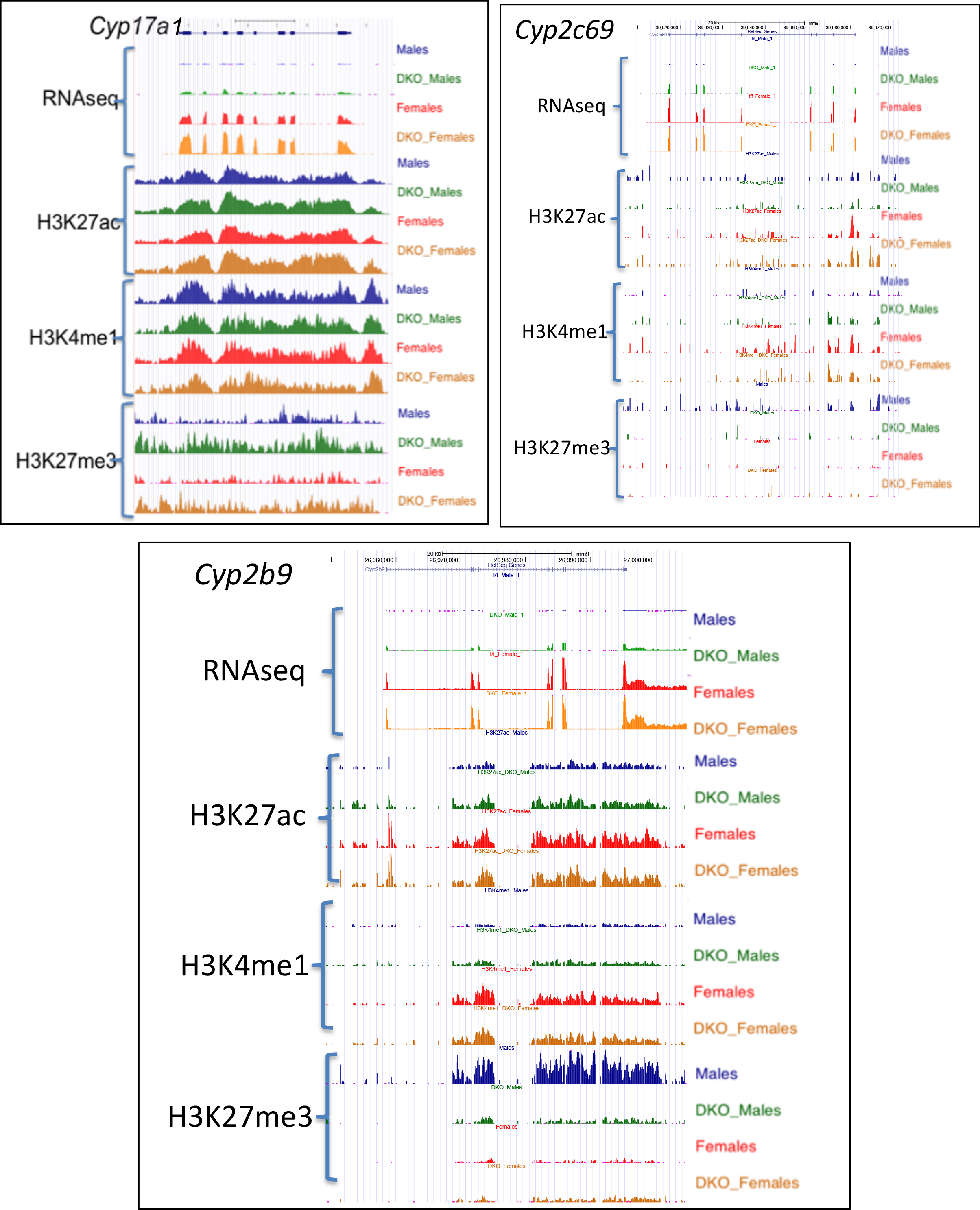
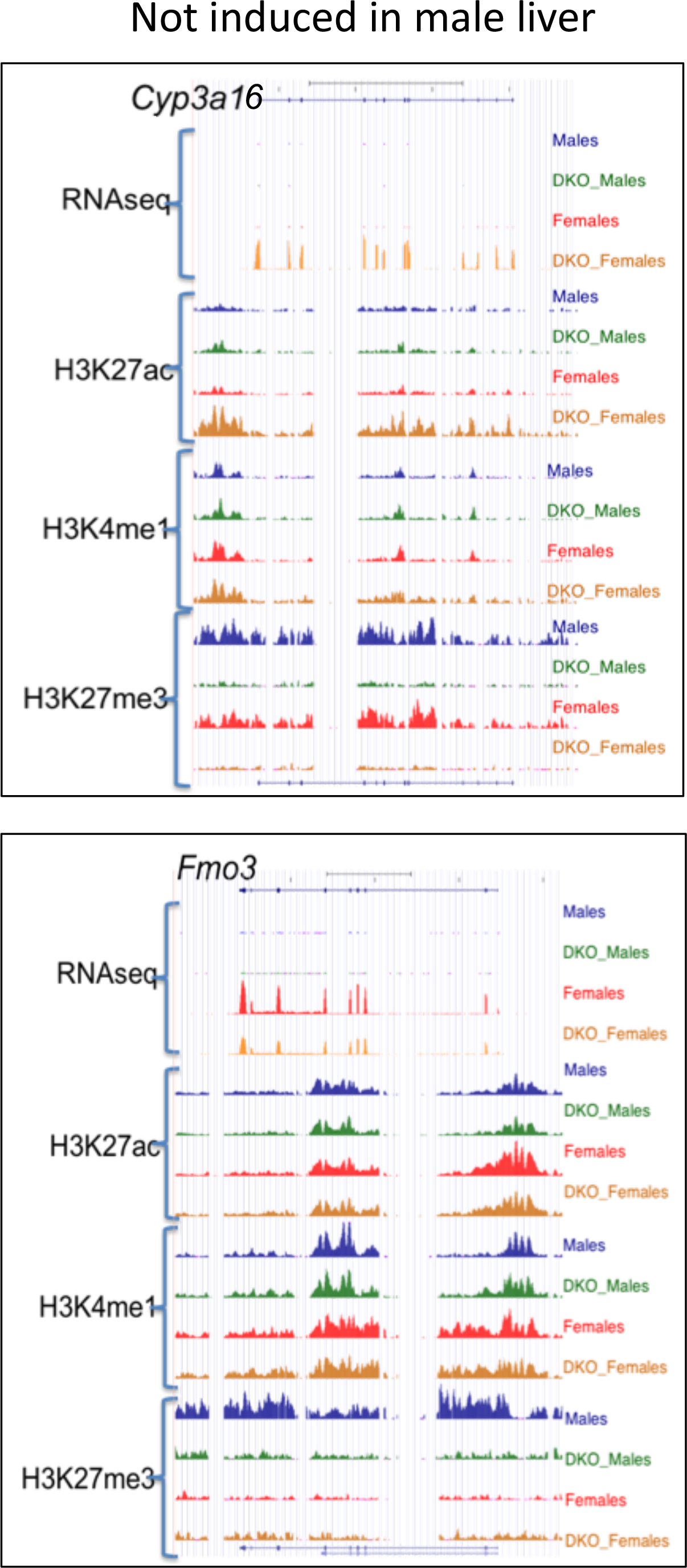
E1/E2-KO-responsive female-biased genes. **A**, top. Listing of expression data for genes whose UCSC Browser screenshots are shown in this figure. Data shown are fold-change and adjusted p-value for each of the following comparisons: Male control vs. Female control, Male E1/E2-KO vs. Male control, Female E1/E2-KO vs. Female control, and Male E1/E2-KO vs Female E1/E2-KO. For those genes that are de-repressed in E1/E2-KO male livers, a feminization percentage and the differential histone mark group is indicated. **A-D**, Shown are UCSC screenshots of normalized RNA-seq or ChIPseq reads for the following E1/E2-KO-responsive female-biased genes: *A1bg, Hao2, Sult2a1, Sult2a5, Sult2a6, Cyp7a1, Cyp2c69, Cyp2b9, Cyp3a16* and *Fmo3.* DKO, Ezh1/Ezh2 double knockout mouse liver.

**Fig. S3.**
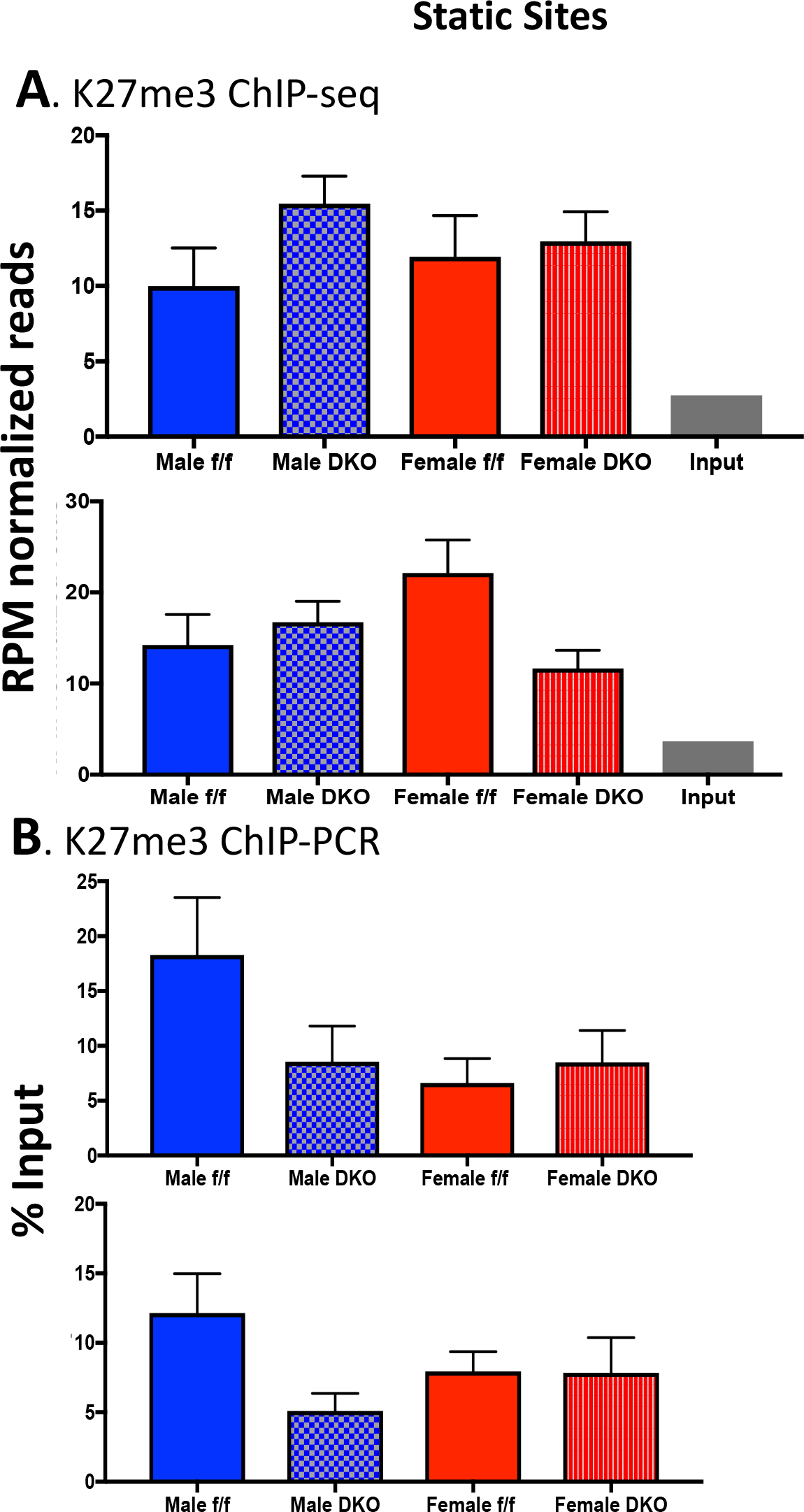
Normalization of H3-K27me3 ChIP-seq. Reads per million (RPM) normalized sequence read counts and ChIP-qPCR validation of static H3K27me3 sites for each of the groups and for the input control. Read counts were obtained for the genomic location corresponding to the qPCR amplicon plus 100 bp to approximate the 200 bp average sequence library insert size. The qPCR data corresponds to % input for n= 4 individuals per group, MEAN +/-SEM. The genomic regions interrogated map to two intergenic static regions (qPCR amplicons: Chr 7: 53,631,859-53,631,958) and Chr 15: 66,117,216-66,117,274). Primers used for qPCR are shown in Table S1B. DKO, Ezh1/Ezh2 double knockout mouse liver.

**Fig. S4.**
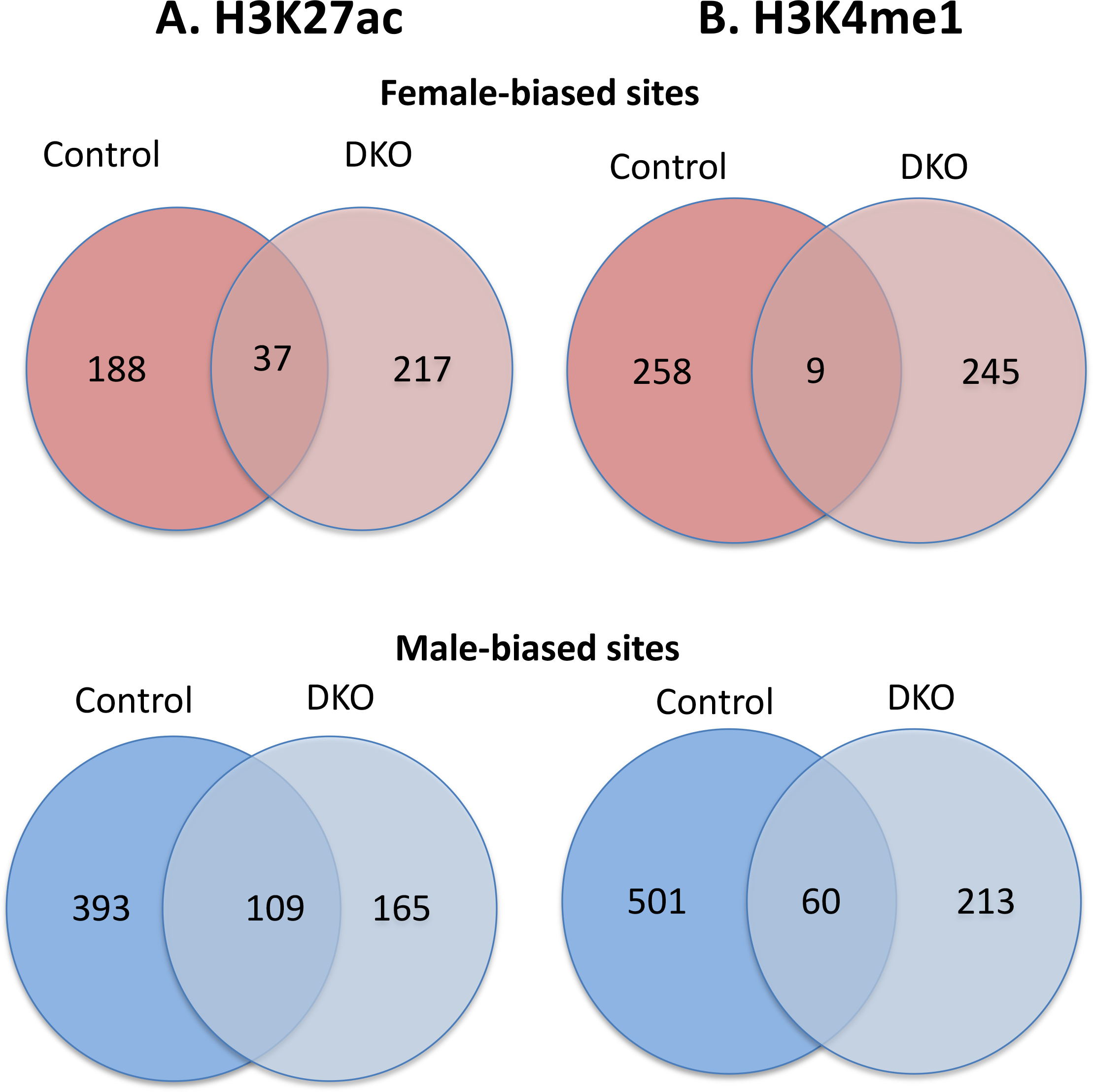
Sex-biased H3K27ac and H3K4me1 sites presented in Fig. 6B. Venn diagrams show the low degree of overlap between the sex-biased sites identified in control livers and those identified in E1/E2-KO livers. This low overlap between the sets of sex-biased H3K27ac and H3K4me1 sites in control, compared to E1/E2-KO mouse liver, indicates that sex-biased chromatin marks are both gained and lost in Ezh1/Ezh2-deficient liver. DKO, Ezh1/Ezh2 double knockout mouse liver.

